# Mating system and environment predict the direction and extent of introgression between incipient *Clarkia* species

**DOI:** 10.1101/2023.08.30.555593

**Authors:** Shelley A. Sianta, David A. Moeller, Yaniv Brandvain

## Abstract

Introgression is pervasive across the tree of life, varying across taxa, geography, and genomes. However, we are only beginning to understand the factors that modulate this variation and how they may be affected by global change. Here, we used 200 genomes and a 15-year site-specific environmental dataset to investigate the effect of mating system divergence and environmental variation on the magnitude of introgression between two recently diverged annual plants. Two subspecies of *Clarkia xantiana* diverged ca. 65k years ago and subsequently came into secondary sympatry where they form replicated contact zones. We found that introgression is asymmetric between taxa, with substantially more introgression from the self-fertilizing taxon to the outcrossing taxon. This asymmetry is caused by a bias in the direction of initial F1 hybrid formation and subsequent backcrossing. We also found extensive variation in the outcrosser’s admixture proportion among contact zones, which is predicted nearly entirely by interannual variance in spring precipitation. Greater fluctuations in spring precipitation result in higher admixture proportions, likely mediated by the effects of spring precipitation on the expression of traits that determine premating reproductive isolation. Climate-driven hybridization dynamics may be particularly affected by global change, potentially reshaping species boundaries and adaptation to novel environments.

## Introduction

Secondary contact provides the ultimate test of whether incipient species have diverged sufficiently to remain distinct. Population genomic analyses have revealed that introgression is quite common for taxa in secondary contact^1–4^ and that introgression varies among contact zones^5–7^ and parts of genomes^1^. However, the ecological factors and phenotypic traits that modulate the direction and magnitude of admixture, and its effects on species boundaries, remain unresolved.

Variation in introgression among independent contact zones may be caused by multiple mechanisms, e.g., differences in relative abundance^6^, the local genetic architecture of reproductive isolation, or the environmental context^7^. The environment can mediate both the expression of traits impacting hybridization^8,9^ and the fitness consequences of hybridization and introgression^10^, making reproductive isolation context dependent. While studies of geographic clines reveal how introgression varies across the species’ range^11–13^, we have limited empirical examples that link specific environmental factors to the extent of introgression in replicated contact zones.

In addition to depending on the environment, the extent of introgression can differ between two hybridizing species. In animals, there is evidence that ancestry from species with more attractive males introgresses into sister taxa with less attractive males than vice versa^14,15^. In plants, too, introgression is often asymmetric. The evolution of self-fertilization from cross-fertilization has occurred repeatedly throughout the evolutionary history of plants and seems to be associated with asymmetric reproductive isolation^16–18^, with apparent introgression more common from selfing to outcrossing lineages^19–24^.

A frontier in speciation genomics is to move beyond tests of whether introgression has occurred and towards prediction of quantitative variation in introgression. Central to this agenda is identifying key drivers that promote hybridization and the persistence of introgressed ancestry. Improved estimates of the magnitude of introgression across many individuals from independent contact zones allows for tests of hypotheses on the causes of variation in reproductive isolation. We examined a well-studied pair of plant taxa, subspecies of *Clarkia xantiana*, that diverged recently^20^, differ in mating system^25^, and form replicated contact zones across a narrow region of secondary sympatry^20,26^. Previous work showed that prezygotic factors cause >98% isolation for both subspecies *xantiana* (outcrosser) and subspecies *parviflora* (selfer)^25^. However, preliminary evidence indicates that introgression is asymmetric, where gene flow is more probable from the selfer to the outcrosser^26^. Finally, the mean and variance in precipitation and temperature, which are the primary drivers of population dynamics^27^, vary strongly across this region of sympatry.

We used 200 whole-genome sequences and 15 years of site-specific weather data to examine the contribution of mating system divergence and environmental variation to the substantial quantitative variation we found in introgression across six independent contact zones occurring over < 30 km. We tested the hypothesis that the magnitude of introgression was greater in more variable environments where opportunities for hybridization may be more frequent. Remarkably, mating system, which causes asymmetry in the direction of hybridization, and variability in spring precipitation, which modulates the frequency of hybridization, nearly fully explain variation in introgression across the contact zones.

## Results

### Genome divergence and population structure

We assembled and annotated a reference genome of an inbred line of the selfer, *parviflora*, using PacBio sequencing (50x) and Bionano optimal maps for scaffolding. The reference genome was assembled into 59 scaffolds totaling 695 Mb, with 45k annotated genes. We then performed whole-genome resequencing (WGS) on 200 individuals from across the range of each taxon (Supplementary Table 1 and Fig. 1a,b). We subset our filtered dataset to coding region sequences and generated two high-quality matrices to use in analyses: all 4-fold degenerate synonymous sites (variant and invariant, 1,751,603 bp) and 4-fold degenerate synonymous biallelic SNPs (353,538 bp).

**Figure 1:**
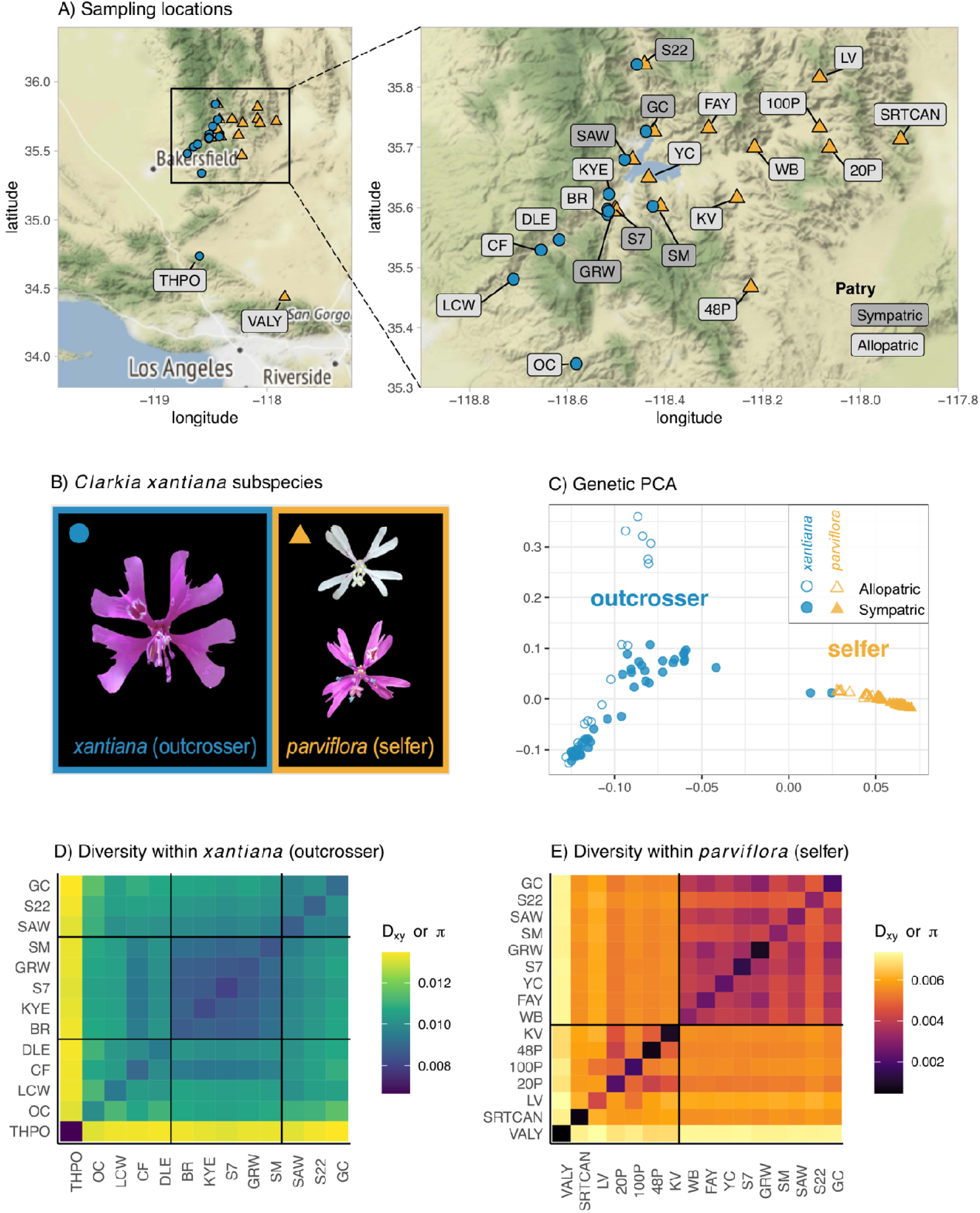
Subspecies of *Clarkia xantiana* are genetically distinct and structured primarily by geography. A) Sampling locations of populations. Sympatric contact zones where both subspecies co-occur are labeled in dark gray, allopatric populations are labeled in light gray. B) Flowers of each subspecies. The selfer has small flowers, varies in flower color, and has less spatial and temporal separation between mature male and female reproductive organs. C) Genetic PCA between subspecies based on 4-fold degenerate SNPs from coding sequences. X-axis is PC1 and y-axis is PC2. Each point represents an individual, with symbol fill reflecting whether that individual is from sympatry or allopatry, respectively. D and E) Genetic diversity within and among populations of each subspecies. Diagonals represent π, diversity within a population. Off-diagonals represent *D*_xy_, divergence between populations. Solid black lines demarcate qualitative groupings of populations based on pairwise D_xy_. D) Diversity within the outcrosser, *xantiana*. E) Diversity within the selfer, *parviflora*.

Genetic PCA showed that the two taxa are genetically distinct (Fig. 1c). Consistent with expectations from their difference in mating system, the selfer, *parviflora*, had one quarter the diversity within populations (π), half of the absolute divergence among populations (*D*_xy_), and four times the relative differentiation among populations (*F*_ST_) relative to the outcrosser, *xantiana* (Table 1 and Extended Data Fig. 1).

**Table 1:** Diversity and divergence within and between species is consistent with mating system shifts. Average (2 * standard error) neutral diversity within populations (π), absolute pairwise divergence among populations (D_xy_), and relative pairwise divergence among populations (F_ST_).

| | $\pi$ | $D_{xy}$ | $F_{ST}$ |
| --- | --- | --- | --- |
| *parviflora* (selfer) | 0.20% (0.06%) | 0.53% (0.02%) | 0.411 (0.03) |
| *xantiana* (outcrosser) | 0.87% (0.05%) | 1.06% (0.03%) | 0.10 (0.02) |
| *parviflora* X *xantiana* | NA | 1.12% (0.01%) | 0.36 (0.01) |

Population structure within each taxon largely mirrored geography. Within the outcrosser, southern populations (THPO, OC, LCW, DLE) were differentiated strongly from northern populations; the latter occur in the region of secondary sympatry and adjacent allopatric populations (Fig. 1d and Extended Data Fig. 2b-d). Within the zone of secondary sympatry, southern outcrosser populations (SM, S7, GRW) were closely related to nearby allopatric outcrosser populations (BR, KYE; Fig. 1d and Extended Data Fig. 2d). By contrast, northern sympatric outcrosser populations (S22, SAW, GC) form a distinct cluster that is divergent from the remaining outcrosser populations (Fig. 1d and Extended Data Fig. 2d). Within the selfer, populations from the region of sympatry and adjacent allopatric sites form a cluster that is strongly divergent from allopatric populations farther from the region of sympatry (Fig. 1e and Extended Data Fig. 2e,f).

### Allopatric origin of selfer with gene flow in secondary sympatry

Prior analyses based on a smaller dataset showed that the selfer diverged from the outcrosser in allopatry ca. 65,000 years bp and later came into secondary contact^20,26^. Consistent with those results, we found that allopatric selfer populations were more similar to the outcrosser on PC1 than sympatric selfer populations (Fig. 1c). Low genetic diversity within and low pairwise *D*_xy_ between sympatric selfer populations (Fig. 1e) suggest that serial bottlenecks accompanied the selfer’s westward range expansion.

Patterns of population structure also supported a model of “budding speciation”, wherein one taxon (here, the selfer) is recently derived from one lineage within another taxon (here, the outcrosser), leading to variation in how diverged different outcrosser populations are from the selfer. Populations of the outcrosser differed dramatically in their divergence from the selfer (median *D*_xy_ ranges from 0.86% to 1.36%), while divergence from the outcrosser was consistent across populations of the selfer (median *D*_xy_ ranges from 1.11% to 1.13%, Fig. 2A). Of all the allopatric outcrosser populations, LCW was the least diverged from the selfer in our dataset (D_xy_ of 1.11%) and can serve as a proxy for the ancestral progenitor population from which the selfer arose. Using LCW as an outgroup, we followed ref.^29^ to estimate a divergence time of ca. 56,000 years bp.

**Figure 2:**
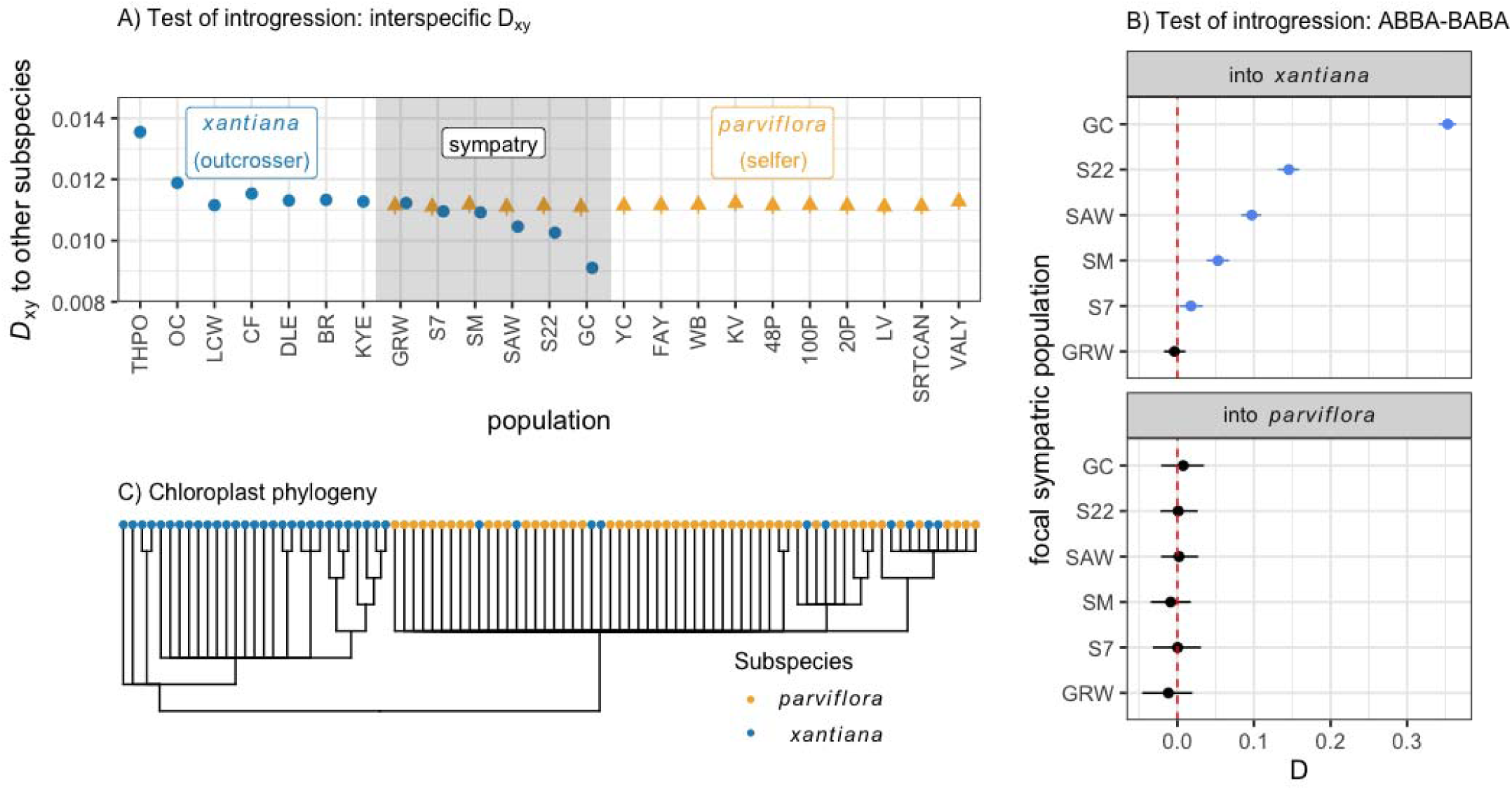
Tests for the presence of introgression find evidence for introgression into the outcrosser, *xantiana*. A) Interspecific D_xy_ between a *xantiana* (blue) or *parviflora* (orange) population at a given site (x-axis) and the other subspecies. Points represent mean pairwise interspecific D_xy_ between the focal population and all other populations of the other subspecies; error bars represent one standard error. Sympatric sites are highlighted by the gray rectangle. B) ABBA-BABA tests of introgression into *xantiana* (top facet) or *parviflora* (bottom facet). Points are the observed D-statistic and error bars represent bootstrapped 95% confidence intervals. Tests of introgression into *xantiana* take the form of (P1= allopatric *xantiana,* P2 = focal sympatric *xantiana* (y-axis), P3 = allopatric *parviflora*, and O = *C. unguiculata*). Tests of introgression into *parviflora* take the form of (P1= allopatric *parviflora,* P2 = focal sympatric *parviflora* (y-axis), P3 = allopatric *xantiana*, and O = *C. unguiculata*). C) Dendrogram of chloroplast phylogeny, with tip points colored by subspecies, shows the 10 *xantiana* individuals that fall within the *parviflora* clade. Seventy percent of individuals within each population are subset out for space; full phylogeny is shown in Extended Data Fig. 4.

Patterns of interspecific divergence (*D*_xy_) are consistent with introgression into the outcrosser but not into the selfer. Interspecific divergence from the selfer is lower for sympatric (mean *D*_xy_ = 1.05%) than for allopatric (mean *D*_xy_ = 1.17%) outcrosser populations (Fig. 2a). This result is robust to the removal of the two southernmost allopatric outcrosser populations (THPO and OC), which are substantially divergent from all other samples (Extended Fig. 2b-d). However, the extent of reduced interspecific *D*_xy_ varied across contact zones. There was apparently little introgression in the southern region of sympatry – *D*_xy_ between the selfer and southern sympatric outcrosser populations (GRW, S7, SM) was comparable to *D*_xy_ between the selfer and nearby allopatric outcrosser populations (Fig. 2a). By contrast, *D*_xy_ between the selfer and northern sympatric outcrosser populations decreased by between 8.4% and 20% relative to average allopatric interspecific *D*_xy_, suggesting extensive introgression in the northern region of sympatry. In contrast, sympatric and allopatric selfer populations had similar *D*_xy_ from the outcrosser (Fig. 2a); this lack of variation in interspecific *D*_xy_ among selfer populations reflects its recent divergence from a taxon with higher diversity (Table 1 and Supplementary Text ‘Issues detecting introgression into a selfer’).

### Introgression is asymmetric and independent across contact zones

We first designed ABBA-BABA tests to formally test for introgression into each taxon at each contact zone. We found evidence for introgression into the outcrosser in all contact zones (significant D-statistic) except for the most southern site, GRW (Fig. 2b and Supplementary Table 2). By contrast, we found no evidence for introgression into the selfer (Fig. 2b). We then designed ABBA-BABA tests to evaluate the extent to which introgression is independent across contact zones by asking whether there is an excess of discordant trees that group the outcrosser and selfer from the same contact zone in a clade (Extended Data Fig. 3a-c). We found evidence for independent introgression occurring in at least the three sympatric sites with substantial introgression (GC, S22 and S7; Extended Data Fig. 3d). By contrast, the remaining sites do not show any signals of significant local introgression, suggesting these contact zones either have shared ancestral introgressed ancestry or lack power to detect independence when introgression is rare.

Phylogenetic analysis of SNPs in the chloroplast resolved a well-supported clade of all outcrosser individuals (95% bootstrap support) that is either sister to (51% bootstrap support) or nested within a clade of all selfer individuals (Fig. 2c and Extended Data Figs. 4 and 5). Within the selfer clade, there were ten outcrosser individuals (based on phenotypes and admixture proportions, see below) from three contact zones (highlighted in Extended Data Fig. 5). Within the outcrosser clade, there were no selfer individuals. This asymmetric chloroplast capture demonstrates that hybrids were formed when the outcrosser sired offspring on the selfer and that hybrids backcrossed to the outcrosser through receipt of pollen.

### Extensive variation in the magnitude of introgression among contact zones

We next quantified individual-level admixture proportions (i.e., percent of the genome that is introgressed) using a Hidden Markov Model (HMM)^30^. We found evidence of introgression into each taxon at all contact zones (Fig. 3 and Supplementary Table 3) that is largely asymmetric, with the outcrosser, *xantiana*, having higher admixture proportions than the selfer, *parviflora*. The asymmetry is particularly stark at contact zones where there are higher overall admixture proportions (S22, GC, SAW, SM), and is not observed in populations with low admixture proportions (S7, GRW).

**Figure 3:**
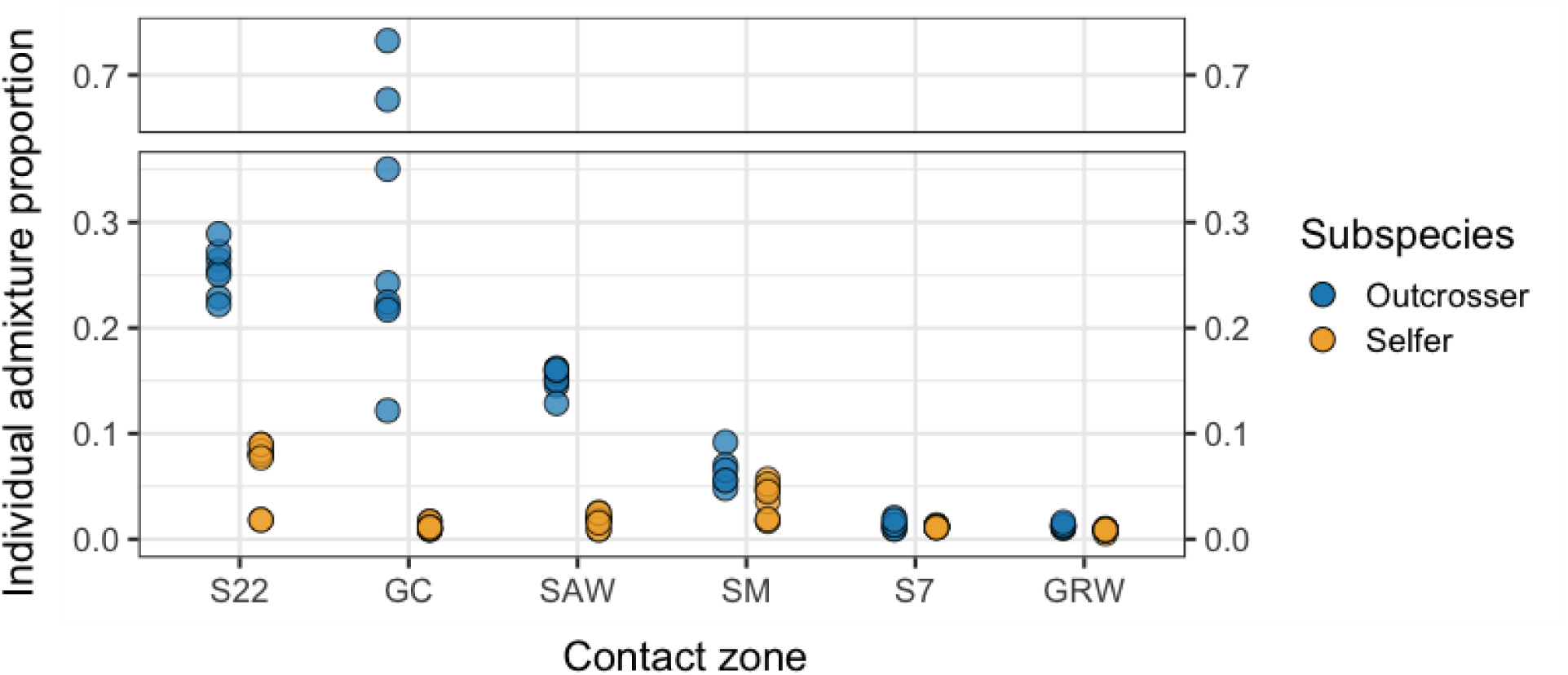
The extent of introgression is largely asymmetric between subspecies and varies across contact zone sites. Hidden Markov Model (HMM) estimates of individual-level admixture proportions. The HMM estimates a probability of ancestry at each site in the genome and individual-level admixture proportions are calculated as proportion heterospecific ancestry, weighted by probability.

Despite ABBA-BABA tests not detecting introgression into the selfer, the HMM identified outcrosser ancestry in the selfer. Admixture proportions were generally low: four populations had admixture proportions under 1.8% and the remaining two populations had average admixture proportions of 3.8% (SM) and 6.9% (S22). Notably, both S22 and SM had the highest sequence diversity (π) and greatest divergence from the other sympatric selfer populations (*D*_xy_, Fig. 1e, Extended Data Fig. 6), consistent with these populations experiencing introgression. We also estimated admixture proportions with the *f*_d_ statistic^31–34^ and detected no introgression into the selfer, but this is likely a byproduct of the low diversity within the selfer (see Discussion, Supplementary Text ‘Issues detecting introgression into the selfer’, ‘Alternative methods to quantifying admixture proportions’). Given the discrepancy among tests, we validated that the HMM truly detected admixed tracks instead of unsorted ancestral polymorphism in the two contact zones with the highest admixture proportions (see Supplementary Text ‘Validation of introgression into the selfer’). We found evidence supporting introgression of the outcrosser into the selfer at S22 (p = 0.029) and marginally significant evidence at site SM (p = 0.075).

The magnitude of introgression into the outcrosser was highly variable among contact zones relative to the selfer. There were high levels of selfer ancestry in outcrosser populations in the northern contact zones (mean admixture proportions of 35%, 25% and 15% in GC, S22, and SAW, respectively), and less introgression into southern contact zones (admixture proportions of 6.5%, 1.4%, and 1.3% in SM, S7 and GRW, respectively). Consistent with observations in our genetic PCA (Fig. 1c), the HMM identified early generation hybrids with exceptionally high admixture proportion at site GC. We also estimated admixture proportions with the *f*_d_ statistic and with a *D*_xy_-based calculation; both methods were highly correlated with admixture proportions estimated by the HMM (see Supplementary Text ‘Alternative methods to quantifying admixture proportions’; Extended Data Fig. 7).

Taken together, the HMM results suggest that introgression was largely asymmetric, with more introgression from the selfer to the outcrosser than the reverse. The HMM also confirmed that introgression dynamics were quite variable across contact zones, particularly for introgression into the outcrosser.

### Greater introgression occurs in more variable environments

Given the exceptional variation in the extent of introgression into the outcrosser across contact zones, we tested whether admixture proportion was related to a set of eight environmental variables collected over 15 years. We included mean and variance of temperature and precipitation in the winter (November - January) and spring (February - May), because natural plants primarily germinate in winter, and primarily grow and flower in spring. Our approach to model selection – leave-one-out (LOO) regression – identified variance in spring precipitation as the strongest predictor of the variation in admixture proportion among outcrosser populations. Variance in spring precipitation was the only variable to outperform an intercept-only model in the LOO test and its performance also exceeded expectations from the null (Permutation-based p = 0.003; Fig. 4a). A simple linear regression similarly identified a strong relationship between variance in spring precipitation and admixture proportion (R^2^ = 0.93; t = 7.02, df = 4, p = 0.002 before; Fig. 4b), with the standardized regression coefficient of 0.96 close to its maximum value of 1.0.

**Figure 4:**
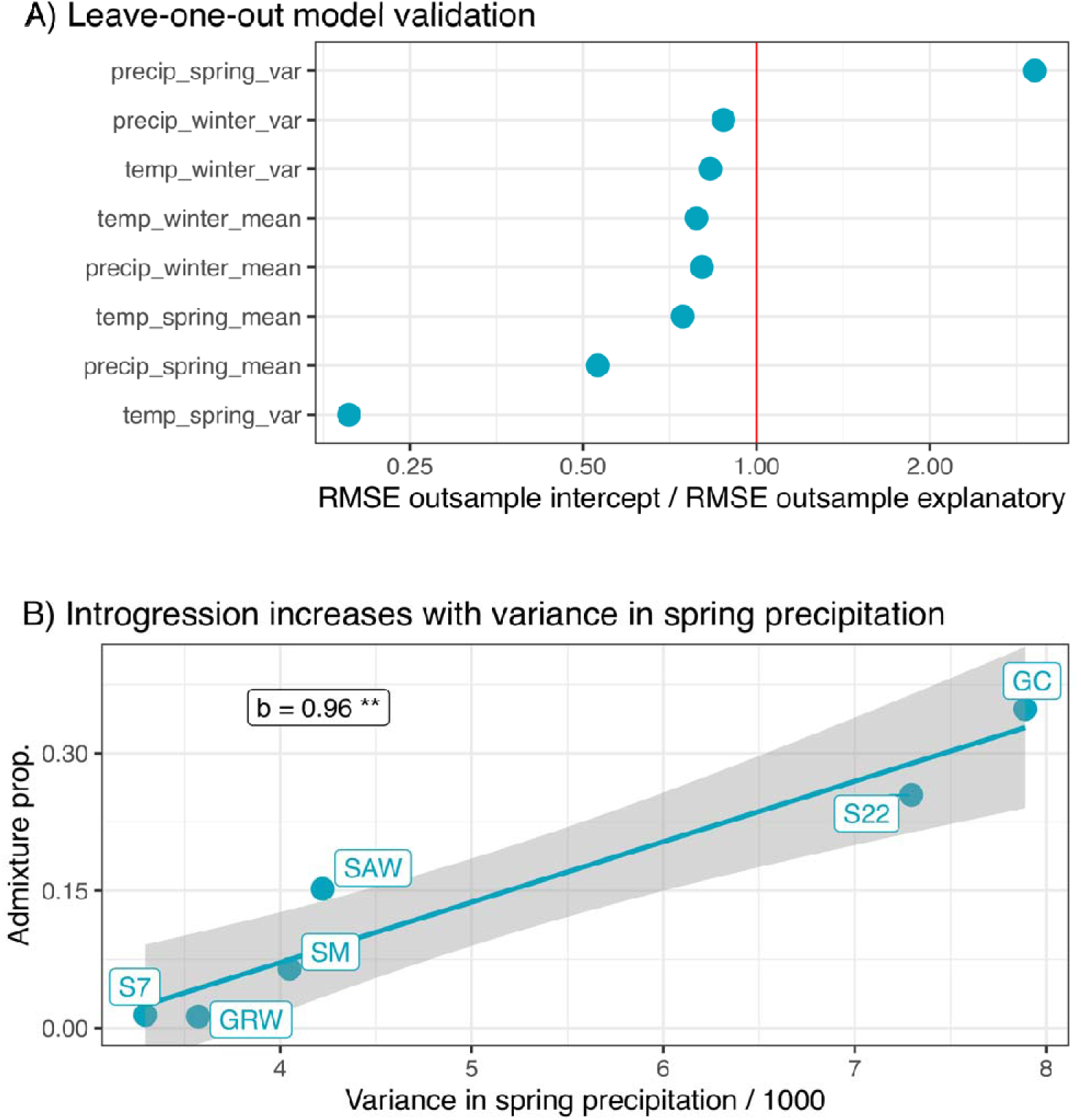
Variance in spring precipitation predicts the admixture proportion across six hybrid zones. A) We present the root mean square error (RMSE) for out-sample prediction from leave-one-out validation for single variable regressions for each of eight environmental variables. To aid in interpretation, we present this as the ratio of outsample RMSE for a model with just the intercept to one with an intercept and slope predicted by the variable of interest (on the y axis). Values exceeding one indicate that the variable of interest predicts admixture proportion in the outsample better than does a simple model with just an intercept. B) The linear relationship between admixture proportion and the variance in spring precipitation. The standardized regression coefficient, *b*, is presented. ** p = 0.002.

## Discussion

Disentangling the factors that modulate the extent to which incipient species remain isolated upon secondary contact is central to understanding the generation of biodiversity. Here, we show that secondary contact between two recently diverged taxa has resulted in extensive variation in the magnitude of introgression, both in the direction between taxa and across six independent contact zones that span a narrow 15 km x 30 km region. Mating system divergence causes asymmetry in introgression, with considerably more introgression from the selfer to the outcrosser than the reverse. Admixture proportion in the outcrosser, which varies strongly across contact zones, is largely explained by the environment, specifically interannual variance in spring precipitation. Environmental fluctuations may result in episodes where reproductive isolation is weakened, thereby increasing the opportunity for gene flow.

While variation in hybridization dynamics and introgression across contact zones have been documented in other systems^10,33^, rarely have studies identified factors modulating that variation (but see refs.^10,34^). In *Clarkia xantiana*, introgression in the outcrosser varies clinally across geography, with higher admixture proportions in northern (range: 15% - 35%) than southern contact zones (range: 1% - 6%). Such variation in admixture across sites likely reflects spatial variation in factors that modulate the opportunity for hybridization (e.g., ref.^10,35^) and/or selection on introgressed ancestry^36–39^. We find one environmental variable– interannual variance in spring precipitation – explains nearly all variation in the outcrosser’s admixture proportion across contact zones.

Variation in spring precipitation is associated with variation in flowering time among closely related species^40,41^. Indeed, divergence in flowering time between the *Clarkia* subspecies is likely driven by divergent adaptation to the onset of late-spring drought: the selfer is a drought-avoider and flowers earlier than the outcrosser, which is a drought-tolerator^41^. This adaptive divergence in flowering time results in strong premating isolation in sympatry – preventing 79-96% of gene flow^42^. However, flowering time variation along environmental gradients is often caused by plasticity, not adaptation^42^; much like adaptive responses, a given environmental variable can lead to differences in the direction and magnitude of flowering time change among species with similar life histories. For example, serpentine soils can induce both earlier and later flowering across 17 annual California wildflowers^43^. As such, when abiotic environments fluctuate through time, the extent of overlap in flowering time may vary in concert because of species’ differing responses to shared environments. Results from this study suggest that *C. xantiana* contact zones with greater variance in spring precipitation may experience greater temporal fluctuations in phenological reproductive isolation, leading to greater opportunities for gene flow and introgression. Past documentation of flowering time in these contact zones, along with 20 years of personal observations, have suggested that flowering times overlap more in years with low spring precipitation (Supplementary Table 4)^25^. Moreover, transplant experiments have shown that flowering times of the two taxa are more similar in some environments than others^17,45^. Taken together, environmental fluctuations may be pivotal for determining the degree of reproductive isolation between incipient species because of its influence on the expression of traits that mediate premating isolation.

Mating system divergence between *xantiana* and parviflora has resulted in asymmetric introgression upon secondary contact. In all contact zones (except two with negligible levels of introgression) there was substantially more introgression into the outcrosser than into the selfer. Our observations of chloroplast variation support the hypothesis that pollen from the outcrosser sires F1 offspring on the selfer –multiple outcrosser individuals from three contact zones were nested within the clade of selfer chloroplast sequences. Moreover, our results suggest that consistent backcrossing to the outcrosser often involves the outcrosser siring offspring on F1 and early generation hybrids. There are several key phenotypic and ecological differences between outcrossers and selfers that likely mediate this direction of gene flow and, ultimately, introgression^17,45–47^. Outcrosses are more abundant^17,46–48^ and they produce considerably more pollen (up to two orders of magnitude)^46^ than selfers, causing the outcrosser pollen pool to be much larger than the selfer pollen pool. In addition, outcrossers are more attractive to bee pollinators^23,24,49–53^ which further increases the probability that transitions occur from outcrossers to selfers. Lastly, outcrosser pollen typically outcompetes selfer pollen when growing in the same style^17,47–49^. In *xantiana/parviflora*, an asymmetric postzygotic barrier (F1 seed viability) in this system also makes it more likely for hybrids to successfully form on a selfer parent^53–55^. Ultimately these factors overwhelm countervailing factors that might increase gene flow in the opposite direction (e.g., self-fertilization preempts ovules from being fertilized by outcrosser pollen). Other studies have found similar asymmetric introgression between selfers and outcrossers^23,24,50–54^, suggesting that these phenotypic and ecological factors associated with mating system shifts have consistent consequences on gene flow and introgression.

Although most studies have not found introgression from an outcrosser into a selfer^50,52,54^ (but see ref.^59^), we detect such introgression, albeit at relatively low levels (< 7%). However, our capacity to detect this direction of introgression depended on the method: genome-wide based methods (e.g., ABBA-BABA, *D*_xy_-based tests of introgression and *f*_d_) did not find evidence of introgression into the selfer. Only the local ancestry analysis, which uses a Hidden Markov Model to detect tracks of ancestry across the genome, detected introgression in this direction. The discrepancy between these types of analyses is likely due to a detection bias in observing introgression into a recently-derived taxon with low diversity, such as into the selfer *parviflora*. The detection bias occurs when divergence between the taxa is comparable to diversity with the outcrosser (as is the case here, Table 1), causing the time to coalescence between an outcrosser and selfer allele to be the same when the selfer allele is introgressed from the outcrosser (coalescence time proportional to diversity within the outcrosser) versus when the selfer allele is not introgressed (coalescence time proportional to interspecific divergence). We advocate for consideration of this detection bias and for extra validation to be performed when investigating introgression in a system with recently diverged taxa that differ in levels of diversity, as may generally be true in cases of budding speciation^55–57^.

Secondary contact in *Clarkia xantiana* has resulted in variable levels of introgression across a small spatial scale, which is nearly fully explained by mating system divergence and spatial variation in interannual spring precipitation. To our knowledge, this is the first study to demonstrate a quantitative link between temporal variance in the environment and introgression. Temporal variance in the environment can cause plastic shifts in traits mediating reproductive isolation (e.g., flowering time) that increase the probability of gene flow. As environments continue to rapidly change, and become more variable, due to climate change, we may expect to see more opportunities for hybridization^58,59^. Reduced reproductive isolation and increased introgression have already been observed in other systems in response to pollution^60^, changes in water turbidity^61,62^, and disturbance^63,64^. An important next step will be to determine whether changes in the rates of introgression with global change have adaptive (e.g., refs^65,66^) or maladaptive consequences in natural settings.

## Methods

### Study system

*Clarkia xantiana* A. Gray (Onagraceae) is a self-compatible, annual plant native to the foothills of the southern Sierra Nevada, Tehachapi and Transverse Mountain Ranges of California, U.S.A^25^. *Clarkia xantiana* is composed of two subspecies – the primarily outcrossing *C. xantiana ssp. xantiana* (hereafter *xantiana*) and the primarily selfing *Clarkia xantiana ssp. parviflora* (hereafter *parviflora*)^73^. Given that these taxa are approximately 99% reproductively isolated^71^, they are likely incipient species as opposed to their original taxonomic description as subspecies. *Xantiana* occurs in the western foothills of the Sierra Nevada in low elevation grasslands, middle elevation oak-pine savanna, and high elevation chapparal. *Parviflora* occurs in low-productivity scrub habitats with occasional pines and Joshua trees. While their ranges are largely allopatric, they overlap in a region centered on Isabella Lake. In their region of overlap the two subspecies co-occur at multiple discrete sites. When sympatric, individuals occur within meters of one another at the boundary of different microhabitats. As such, they have parapatric distributions at both fine and geographic scales.

### Constructing the first *Clarkia* reference genome

We assembled and annotated the first *Clarkia* reference genome. The genome was derived from one *parviflora* (selfer) individual that was selfed for five generations. We chose this individual from the Long Valley population (LV on Fig. 1a) because prior analyses indicated that this was the earliest diverging, extant *parviflora* population^26^, and thus should minimize reference bias when mapping sequences from *parviflora* and *xantiana.* We combined PacBio sequencing and Bionano optical mapping to generate a 695.6 Mbp genome assembled on 59 scaffolds with an N50 length of 32.5 Mbp. We created a genome annotation by combining gene predictions from transcriptome data (from leaf, root, stem, flower, and fruit tissue) and a suite of ab-initio gene predictor programs using the funannotate pipeline (1.8.9^74^), resulting in 44,998 total gene models. We assembled a reference genome for the chloroplast by aligning raw PacBio reads to the chloroplast genome of a species from a closely-related genus (*Oenothera villarica*, Onagraceae; NCBI accession GI 1034702878). We used Canu (v 2.1)^75^ to assemble aligned raw reads, resulting in one continuous contig 161,015 nucleotides long. Full details about how the reference genome was assembled and annotated are provided in the Supplementary Methods under ‘Reference Genome’.

### Whole-genome-resequencing of the outcrosser (*xantiana*) and selfer (*parviflora*)

To quantify the distribution of genetic diversity within and between subspecies, and examine patterns of introgression in the region of secondary sympatry, we collected range-wide samples of each subspecies (Fig. 1a; Supplementary Table 1). In the summer of 2019, we collected seeds from 13 *xantiana* populations (six in contact zones and seven allopatric) and 16 *parviflora* populations (six in contact zones and ten allopatric). We also collected seeds from four congeneric species for use as outgroups in analyses (Supplementary Table 1). We germinated field-collected seeds and collected leaf tissue for whole-genome-resequencing. We sampled an average of 8.5 and 5 individuals from contact zone and allopatric populations, respectively. In total, we sequenced 200 individuals.

DNA was extracted using the CTAB method and libraries were prepared with the Illumina TruSeq NanoDNA library preparation kit. Samples were split into two unique sequencing pools, each containing 100 samples. Each pool was sequenced across two lanes of an Illumina NovaSeq S4 2×150-bp run, with a targeted depth of 13x.

We removed adaptor sequences and low quality reads with scythe^76^. We used BWA mem to map trimmed reads to the masked reference genome. We used Samtools^75^ to sort reads, Picard to add read groups, and Samtools to index alignments. We used GATK (v 3.8.0)^77^ to call genotypes. GATK’s HaplotypeCaller function was used to generate per-individual genotypes, which were subsequently grouped for joint genotyping with the ‘GenotypeGVCF’ function. Because joint genotyping with GenotypeGVCF assumes random mating among samples, we grouped our samples differently for the outcrosser, *xantiana*, and the selfer, *parviflora*. For the selfer, we did joint genotyping for only individuals within a population because there is usually high differentiation among populations (F_ST_) in selfing taxa. We grouped all samples of the outcrosser, *xantiana,* together for joint genotyping, except for individuals from the southernmost population THPO, which is geographically disjunct from the other samples. The three individuals from each of the four outgroup species were grouped together for joint genotyping. We ran GenotypeGVCF over each scaffold and merged all VCFs for a given scaffold (i.e., from the different groups) into one VCF.

We filtered our VCFs in multiple steps. We first surveyed average depth and percent missing data across all individuals and scaffolds using the BCFtools^78^ stat function and custom R scripts. We flagged 7 scaffolds for which the vast majority of individuals had low (< 5) or high (> 30) depth; these same 7 scaffolds were flagged for having the vast majority of individuals with > 75% missing data. We excluded these seven scaffolds, eliminating 1.2% of the genome. We also flagged and removed 13 individuals that had > 50% missing data across the genome. Paralogs can create mapping issues by having multiple paralogs map to the same regions. We identified and removed regions with potential paralogs by identifying regions with high heterozygosity in the selfer. We flagged sites at which >= 50 of 100 selfer individuals were heterozygous (hereafter high heterozygosity sites), binned high heterozygosity sites into 1kb windows, and removed windows for which there were > 15 high heterozygosity sites. We then filtered out all sites with ‘LowQual’ scores in the FILTER column and changed all genotypes to missing for any individual/site combination where read depth < 4 or > 30. We then removed any site that was missing data for over 50% of the outcrosser individuals and 50% of the selfer individuals. After these filtering steps, we reduced the genome by an additional 73%, resulting in 171Mb.

To ensure that we were using high quality sites for analyses, we further filtered our dataset to 4-fold degenerate sites in coding sequences and to biallelic SNPs within that dataset.

### Quantifying genetic diversity within and between species

We first compared the extent of genetic divergence within and between the two subspecies using PCA. We calculated a genomic PCA using a custom R script to account for missing genotype data by calculating a pairwise genotypic covariance matrix across SNPs at fourfold degenerate sites and finding components as the eigenvectors of this matrix. To better visualize population structure within species, we generated intra-subspecific PCAs as above, discarding geographically distant outlier populations from the southern edge of the range for each subspecies (VALY in the selfer, and OC and THPO in the out), which had undue influence on PC space (see Extended Data Fig. 2).

To determine how genetic variation is structured across the range of each subspecies, we characterized the mean number of pairwise sequence differences at fourfold degenerate sites (an estimate of π within populations, and *D*_xy_ between populations), and population differentiation (measured as *F*_ST_) with the program, pixy^29^.

We asked whether patterns of interspecific *D*_xy_ were consistent with an allopatric origin of the selfer, *parviflora*, wherein the selfer arose from a historic outcrosser population that was more closely related to one of the allopatric (southern) outcrosser populations than one of the sympatric (northern) populations, as proposed by ref.^79^. We found that the allopatric outcrosser population LCW was the least diverged with the selfer. We used LCW as a proxy for the ancestral outcrosser population from which the selfer arose to estimate a population split time. We did this with the method described in ref.^80^ that uses the difference between divergence between taxa and diversity within this (proxy for the) ancestral population divided by two times the mutation rate (using the standard assumption of 1.5 x 10^-8^ mutations per base pair mutation per generation) to estimate a population split time.

We also used interspecific *D*_xy_ to determine if there is evidence for gene flow in sympatry. We expect gene flow to decrease divergence (interspecific *D*_xy_) between a sympatric outcrosser population with selfer ancestry and all selfer populations, but such gene flow would not affect divergence between the source selfer populations and allopatric outcrosser populations. In principle, the same logic should apply for cases of gene flow from the outcrosser to the selfer. However, because divergence between outcrosser populations is comparable to divergence between the subspecies, this approach is underpowered to detect gene flow in this direction (see Supplementary Text ‘Issues detecting introgression into a selfer’).

### Tests of introgression

We formally tested for the presence of introgression by designing ABBA-BABA tests^27^ for introgression into each subspecies at each sympatric site. In ABBA-BABA tests, the four taxa are labeled P1, P2, P3, and O for outgroup. P1 and P2 are populations of the hypothesized recipient species, where P1 does not experience introgression but P2 does. P3 is a population of the hypothesized donor species. Among discordant trees that do not match population history, introgression results in more (P2,P3)(P1,P4) trees than (P1,P3)(P2,P4) trees, while no such excess is predicted without introgression. We conducted 12 such ABBA-BABA tests – in each test, one of the six contact zone populations of each subspecies was designated as the putative recipient of gene flow, P2, and the outgroup was set as the congener, C. *unguiculata,* in all tests. For tests of introgression into the outcrosser, P1 was the allopatric outcrosser population DLE and P3 was the allopatric selfer population FAY. For tests of introgression into the selfer, we swapped P1and P3. By using allopatric populations as P1 and P3, we can separately test for introgression in both directions. For all tests, we calculated the number of discordant trees across the genome at coding sequence sites, calculated the imbalance in discordant tree topologies using the D-statistic, and evaluated significance based on 1000 bootstrap replicates and whether the bootstrapped 95% confidence intervals overlapped zero.

We then tested the null hypothesis that the observed introgression across contact zones represents one historical admixture event in which introgressed ancestry spread across the region of sympatry or independent occurrences of introgression at each contact zone (or some combination of the two). To do so, we used 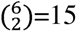 ABBA-BABA tests, each of which consisted of outcrosser and selfer samples from two contact zones. We tested for an excess of discordant trees that grouped populations by contact zone site (Extended Data Fig. 3, tree cartoon, A-C).

We investigated patterns of relatedness among chloroplast sequences of the two taxa to determine if there was a consistent direction of gene flow occurring between the taxa. For example, if one subspecies always acted as the seed parent in the F1 cross and backcrossing consistently occurred though pollen flow of the other subspecies, we would expect to find the donor species chloroplast in the recipient species. We extracted the chloroplast genome, using the coordinates identified when blasting our chloroplast assembly to the reference genome, from our whole-genome-resequencing dataset. We removed the 13 individuals that had high missing data (as above), removed LowQuality sites from the FILTER column of the VCF, turned genotypes with sample depth < 10 to missing, removed sites with > 20% missing data for either subspecies and subsetted to only biallelic SNPs. We converted our filtered VCF to a fasta (vcf2phylip.py script from https://github.com/edgardomortiz/vcf2phylip) and inferred the phylogenetic relationships among taxa with RAxML version 8.2.11^80^ using the GTR model and rapid bootstrapping. We used the four congeneric *Clarkia* species (Supplementary Table 1) as outgroups.

### Quantifying admixture proportions across contact zone sites

In addition to testing for the presence and independence of introgression, we wanted to quantify the extent to which introgression was occurring in the two subspecies across the region of sympatry. We used a Hidden Markov Model (HMM) to infer ancestry across the genome and to estimate the genome-wide admixture proportion for each individual^29^. These models use a reference panel containing allele frequencies at every site for each subspecies and a transition matrix based on recombination rates to estimate posterior distributions of three ancestry genotypes at each site: homozygous for the conspecific ancestry, heterozygous for both ancestry types or homozygous for the heterospecific ancestry. To construct reference panels for each subspecies, we selected individuals from three allopatric populations of each taxon that were spatially proximate to the contact zone. We used FAY, KV, and WB for the selfer and CF, DLE, and KYE for the outcrosser. We ran the ancestry HMM on each subspecies/contact zone combination separately, for both subspecies in the six contact zones, using a SNP matrix of 4-fold synonymous coding region sites.

To parameterize the emission probabilities, ancestry HMM requires an estimate of global (population-level) admixture proportions in the sample. As we did not *a priori* know this value, we initially assumed it to be 10% and then used the Baum-Welch expectation-maximization algorithm to update this estimate, as suggested by ref.^29^. We reran the HMM until the output and input admixture proportions differed by less than 0.001. Because we do not have a high-resolution genetic map for *Clarkia xantiana* we assumed a uniform recombination rate of 1 x 10^-8^ morgans/ bp.

At the end of the final HMM run for each population of each subspecies, we calculate individual admixture proportions in two ways. First, we used only sites that had high confidence ancestry calls (posterior probability of ancestry > 0.8). Second, we used all sites, with ancestry calls weighted by their posterior probabilities. Both estimates are highly correlated (r = 0.9; Extended Data Fig. 7), and we present the estimates using all sites in the main text.

Because we find introgression into the selfer with the HMM local ancestry analysis but not with genome-wide ABBA-BABA analyses, we confirmed that the HMM was inferring introgression versus unsorted ancestral polymorphism using *D*_xy_-based logic (see Supplementary Methods ‘Validation of introgression into the selfer’). We compared our results from the HMM with two additional estimates of the admixture proportion – the *f_d_* statistic ^30^ and a D_xy_-based calculation (see Supplementary Methods ‘Alternative methods to quantifying admixture proportions’). Because all estimates were strongly correlated (Extended Data Fig. 7 and Supplementary Fig. 6) we only discuss results from the HMM in the main text.

### Explaining variation in admixture proportion among populations

We found substantial variation in the magnitude of introgression into the outcrosser, *xantiana*, across contact zones. Guided by these results, we tested whether the outcrosser’s admixture proportion among the six contact zones was correlated with a set of eight environmental variables. These environmental variables were collected over 15 years at each contact zone except S7. For site S7, we used weather station data from a spatially proximate (1.1 km) *xantiana* population with the same slope aspect, BR. The variables we included in our models were mean and variance of temperature and precipitation in the winter (November - January) and spring (February - May), because natural plants primarily germinate in winter and primarily grow and flower in spring. Prior demographic studies have also shown that environmental variance can be pivotal in determining population dynamics ^27^. To overcome the challenges of modeling fewer observed responses (n = 6) than potential explanatory variables (n = 8), we used a leave-one-out (LOO) validation method to select an appropriate model. The LOO validation compares eight regression models (one for each environmental variable) to a model with just the intercept. For each environmental variable, we built six linear models, each one “holding out” one data point. We then summarized model performance for a given environmental variable as the square root of the mean squared difference between the model’s prediction for the missing data point and its observed value (i.e., the Root Mean Squared Error, RMSE). An RMSE exceeding that of the “intercept only” model means that the environmental variable can predict the admixture proportion. We report p-values from both a simple linear regression of our best model, and from a permutation comparing the observed RMSE from leave-one-out validation to its distribution under the 720 ways we could permute the order of the six observations.

We estimated the effect of our environmental variable on admixture proportions by finding the standardized regression coefficient (i.e., the regression coefficient when X and Y are Z-transformed). The standardized regression coefficient can take values from −1 to 1, and by convention values between 0.3 and 0.5 are considered modest effects, and values greater than 0.5 are considered strong effects.

## Acknowledgements

We thank V. Eckhart for collecting the weather station data and A. Raduski for help with genome assembly and calling whole-genome-sequence genotypes. We thank the University of Minnesota Genomics Center for their guidance and for performing DNA extraction, library preparation, and sequencing. The Minnesota Supercomputing Institute (MSI) at the University of Minnesota provided resources that contributed to the research results reported within this article. Funding for this project was provided by NSF award DEB-1754246 to DAM and YB.

## Author Contributions

YB and DAM designed the project. DAM collected samples. SAS and YB analyzed data. SAS wrote the first manuscript and all authors contributed to the final version.

## Data Availability

All genome and resequencing data (PacBio reads for genome sequence, genome assembly, RNA-seq, whole-genome-resequencing data) have been submitted to NCBI (BioProject number PRJNA906640). All other data reported in this manuscript are archived in Figshare (10.6084/m9.figshare.23961792).

## Code Availability

The code used for this manuscript is archived in Figshare (10.6084/m9.figshare.23961792).

