## Supplementary Information for "Mating system and environment predict the direction and extent of introgression between incipient *Clarkia* species"

Shelley A. Sianta *et al.*

Content:

1. Supplementary Text (Methods and Results)

2. Supplementary Figures

3. Supplementary Tables

### **Supplementary Text (Methods and Results)**

#### **Reference genome**

***Sampling and sequencing***

We sequenced an inbred line derived from one *parviflora* individual from the Long Valley population (LV on Figure 1A) that was selfed for five generations. We chose Long Valley because prior analyses indicated that this was the earliest diverging, extant *parviflora* population^1^, and thus should minimize reference bias when mapping sequences from *parviflora* and *xantiana*. We collected young leaf tissue from an individual that was dark treated for 48 hours and extracted DNA with the Nucleon PhytoPure DNA Extraction kit (Cytiva RPN-8511).

Libraries were constructed with the PacBio SMRTbell Express Template Preparation Kit (2.0), removing inserts shorter than 30 kb, and genomic DNA was sequenced by the University of Minnesota Genomics Center (UMGC) on 6 SMRT cells on the PacBio Sequel platform (the 3.0 chemistry). We obtained 60 GB of raw sequence data (approximately 60x genome coverage), with a mean +/- SE longest subread length of 13.1 +/- 0.6 kb and a mean +/- SE longest subread N50 of 21.3 +/- 0.8 kb.

***Assembly***

PacBio reads were assembled using Canu (version 1.9)^2,3^. We ran Canu with an error correction rate of 4.5% and an estimated genome size of 1 GB. After read trimming and correction, our assembly consisted of 884 contigs with an average depth of 36x. BUSCO analysis found 91.2% complete BUSCOs (66.1% single-copy and 25.1% duplicated BUSCOs), 1.9% fragmented BUSCOs, and 6.9% missing BUSCOs.

We used BioNano optical mapping (Corteva Agriscience) to create a backbone on which to scaffold PacBio contigs. High molecular weight DNA was extracted from the same inbred line used for PacBio sequencing. The BioNano genome map contained 93 scaffolds with a total length of 782.1 Mbp and a N50 length of 32.4 Mbp. Combining the BioNano genome map with the PacBio Canu assembly resulted in a hybrid scaffold consisting of 59 primary scaffolds that covered 695.6 Mpb (87% of the PacBio assembly) with an N50 length of 32.5 Mbp. These 59 primary scaffolds combined 79 of the PacBio contigs; the remaining 804 contigs from the PacBio sequence were not placed onto the BioNano genome map. In subsequent analyses, we only use the 59 hybrid scaffolds.

***Annotation***

We used a mix of gene model predictors and transcriptome data to annotate the genome. For the transcriptome, we extracted RNA from a mix of leaf, root, stem, flower and fruit tissue from both well-watered and drought-stressed replicates of the inbred *parviflora* line. RNA was extracted with the RNeasy Power Plant Kit (Qiagen), samples were pooled, and a single library was constructed with the TruSeq Stranded Total RNA workflow with plant RiboZero reduction. The library was sequenced over four lanes of an Illumina NextSeq Mid-output 2x150-bp run, generating 149M read pairs with an average of 89% of bases in each lane having above a Q30 quality score. We removed adaptor sequences and low quality reads with scythe (version 0.994 BETA)^4^.

We used the program funannotate (1.8.9)^5^ a wrapper that uses an array of annotation software, to generate gene prediction models with multiple sources of evidence. We first soft-masked our genome using RepeatModeler (2.0.1)^6^ and RepeatMasker (4.0.5)^7^. RepeatModeler was run on the genome to build a custom library of repeats (TE families) for our genome; 1995 TE families were identified. RepeakMasker soft-masked 68.21% of the genome: half of the masked repeats were identified as retroelements (primarily LTR elements) and the other half were unclassified repeats. The funannotate *train* command generated 32,843 gene predictions from transcriptome data, using Trinity (2.8.5)^8^ and PASA (2.3.3)^9^ to assemble and map transcripts. In the funannotate *predict* command, a suite of ab-initio gene predictor programs (Augustus^10^, SNAP^11^, Glimmerhmm^12^, Genemark^13^) were trained with the PASA gene predictions and a final program, EvidenceModeler^9^, combined all gene predictions. The final annotation consisted of 44,998 total gene models.

***Chloroplast genome***

To assemble a chloroplast reference genome from our PacBio data, we first aligned raw PacBio reads to the chloroplast genome of a species from a closely-related genus (*Oenothera villarica*, Onagraceae; NCBI accession GI 1034702878) using BLASR^14^. We used Samtools (v1.14)^15^ to convert .sam files to .bam files, merge .bam files and convert the merged .bam file to a .fastq file. We used the program Canu (v 2.1)^3,16^ to assemble aligned raw reads, resulting in one continuous contig 161,015 nucleotides long. We then mapped the Canu chloroplast assembly to the BioNano scaffolded assembly. The Canu assembly mapped to one stretch of the BioNano scaffold 100056, comprising approximately 27% of the scaffold.

#### **Issues detecting introgression into a selfer**

##### ***D_xy_-based inference of introgression***

The logic of the *D*_xy_-based test for introgression is that introgression from one taxon to another should lower interspecific *D*_xy_ between the two taxa compared to a case with no introgression. This approach can identify the direction of introgression by comparing divergence of a putatively admixed focal (sympatric) population of one taxon to a putatively unadmixed (allopatric) population of the other taxon. We did not see a decrease in interspecific *D*_xy_ in sympatric populations of the selfer, *parviflora*, relative to allopatric selfer populations (Main Text Fig. 2a), as would be expected if there was introgression in sympatry. However, the lack of a decrease in *D*_xy_ is partially expected, even if the selfer is the recipient of introgression.

This is because the genome-wide average *D*_xy_ of a genome that has introgressed DNA can be broken down into two types of regions: 1) unadmixed regions characterized by the average number of differences between genomes of the two taxa (*D*_xy_ between species), and 2) admixed regions characterized by the average number of differences between genomes of the taxon that donated the introgressed DNA (π within the donor species). When π within the donor species is much lower than *D*_xy_ between the species, introgression results in a lowering of genome-wide *D*_xy_. However, as in the case of the selfer-outcross pair in this study, when π within the donor (outcrosser) species is comparable to *D*_xy_ between the species (Main text Table 2), introgression results in very small reductions in genome-wide *D*_xy_ in the selfer.

As an example, we conduct a brief “back-of-the-envelope” comparison of the strength of signal in *D*_xy_ reduction with symmetric admixture proportions of 20% in both *parviflora* and *xantiana* S7 populations (a sympatric site with little evidence for introgression in either direction). So if 20% of the S7 *xantiana* genome is recently derived from S7 *parviflora*, *D*_xy_ between S7 *xantiana*, and a close allopatric *parviflora* population (YC) would be

= 0.80 * *D*_(X_S7, P_YC)_ + 0.20 * *D*_(P_S7, P_YC)_

= 0.80 * 0.01096550 + -.20 * 0.00382856

= 0.009538

This represents an approximately 13% decrease in *D*_xy_ relative to empirical observations.

By contrast if 20% of the S7 *parviflora* genome is recently derived from S7 *xantiana*, *D*_xy_ between S7 *parviflora*, and a close allopatric *xantiana* population (BR) would be __.

= 0.80 * *D*_(X_S7, P_YC)_ + 0.20 * *D*_(X_S7, X_BR)_

= 0.80 * 0.01096550 + 0.20 * 0.00869099

= 0.0105106

This represents an approximately 4% decrease in *D*_xy_ relative to empirical observations. In fact, observing a reduction in *D*_xy_ comparable to what is observed with 30% introgression in the other direction requires that nearly 60% of the S7 parviflora genome must be recently derived from the S7 xantiana population.

##### ***ABBA-BABA tests***

We similarly failed to detect introgression into the selfer with genome-wide ABBA-BABA tests. We argue that this is due to low power, and an issue for detecting introgression into any species with relatively little genetic variation. For example, in our ABBA-BABA tests where P1 and P2 are *parviflora*, and P3 is *xantiana,* we expect the low diversity in *parviflora* to result in most trees following the species tree (P1 and P2 as sister taxa). This lowers the power available for calculating the D-statistic, which compares the imbalance of two types of discordant gene trees (P1 and P3 vs. P2 and P3 as sister taxa). Indeed, tests for introgression into *parviflora* have many more SNPs following the species tree (BBAA) relative to discordant trees (ABBA, BABA) than do tests for introgression into *xantiana* (Supplementary Table 2).

#### **Validation of introgression into the selfer**

##### ***Rationale***

Because the HMM identified introgression from the outcrosser, *xantiana*, to the selfer, *parviflora*, but neither the interspecific *D*_xy_ nor ABBA-BABA tests found evidence for introgression in this direction, we tested the null hypothesis that introgression uncovered by the HMM reflects mistakes in the HMM rather than actual introgression. The HMM uses allele frequencies of two reference panels - one for each subspecies - to infer ancestry. Given the recent divergence of the selfer from the outcrosser, and drift in the selfer, it may be the case that selfer individuals in the contact zones have different allele frequencies relative to the selfer reference panel because of unsorted ancestral polymorphism, and that the HMM would misclassify genomic regions with such unusual allele frequencies as “admixed”.

To distinguish between true introgression and mistakes in the HMM, we compare patterns of interspecific sequence divergence in genomic regions of the selfer called “admixed” or “unadmixed” by the HMM. If unusual patterns of unsorted ancestral polymorphism are tricking the HMM into wrongly inferring a region is introgressed, we do not expect regions called “admixed” by the HMM to have lower *D*_xy_ with the outcrosser than comparable regions called “unadmixed.” However, if the HMM is identifying truly introgressed genomic regions, we expected “admixed” regions to have lower *D*_xy_ with the outcrosser than comparable “unadmixed” regions.

For every genomic window that is polymorphic for ancestry calls within a given selfer population, we randomly choose one individual with an “admixed” HMM call and one individual with a “conspecific” (or “unadmixed”) HMM call. For each window we calculated *D*_xy_ between an allopatric xantiana population, BR, and the random admixed and random unadmixed sample. We used a permutation test to evaluate the null hypothesis that *D*_xy_ between BR and putatively admixed regions was not lower than *D*_xy_ between BR and putatively unadmixed regions.

##### ***Methods***

The validation method below was performed in each of the two selfing populations for which the HMM detected substantial introgression: S22 and SM. Details of the method are depicted in Supplementary Fig. 1.

We assigned all sites used in the HMM to a 5kb window and called one genotype (homozygous for the conspecific ancestry, homozygous for the heterospecific ancestry, or heterozygous for the two ancestry types) per window, depending on which genotype call had the highest posterior probability. We filtered out sites where the most-confident genotype call had a posterior probability less than 0.9. For each window, we randomly selected one individual within the selfing population that was homozygous for the conspecific ancestry (“conspecific”) and one individual that was homozygous for the heterospecific ancestry (“admixed”), and focused our analysis exclusively on genomic windows polymorphic for ancestry in the focal selfing population.

This resulted in a paired design - for every window there was a chosen individual with a HMM call of conspecific and an individual with a HMM call of admixed. We chose this approach of comparing the same genomic region in admixed and un-admixed regions at the same genomic location from individuals from the same population to remove all other sources of differences in sequence divergence.

We then created two BED files per individual - one with all windows for which that individual was chosen as the representative for introgressed ancestry in the comparison, and one with all windows for which that individual was chosen as the representative conspecific HMM call in the comparison. We used these BED files in the program pixy (Korunes and Samuk 2021) to calculate pairwise *D*_xy_ in windows between the selfer individual and individuals from the outcrosser population BR. We chose this outcrosser population because it is closely related to the southern contact zone outcrosser populations but is allopatric and thus presumably has no introgression. We concatenated all pixy outputs, such that the final pixy output contained, for each window and each HMM call (“admixed”/“conspecific”), pairwise *D*_xy_ between the chosen *parviflora* (selfer) individual and each individual from the outcrosser (*xantiana*) BR population.

We permuted the data to test whether *D*_xy_ between the selfer and outcrosser was significantly less in windows with admixed HMM calls vs conspecific HMM calls. Because the pixy output contained multiple *D*_xy_ values per window and HMM call combination, causing pseudoreplication, we combined permutations with subsampling. For each permutation step, we randomly selected one *D*_xy_ value from each window/HMM-call combination. We calculated the observed difference in genome-wide *D*_xy_ between admixed and conspecific calls. We then permuted the HMM call values across the subsampled dataset and recalculated the difference in genome-wide *D*_xy_ between admixed and conspecific calls. We also took the difference between the observed difference and the permuted difference. We repeated this permutation step 1000 times to build distributions of the observed difference and the permuted difference in *D*_xy_ between admixed and conspecific calls. We then calculated a p-value as the proportion cases in which the trued observation exceeded the permuted value.

##### ***Results***

We find evidence in the selfer, *parviflora*, populations S22 and SM that sites identified as admixed by the HMM are truly due to introgression from the outcrosser, *xantiana*, and not from unsorted ancestral polymorphism.

At S22 we found that there was a significant increase in genome-wide *D*_xy_ between the outcrosser and the selfer when genomic windows were characterized by conspecific HMM calls (average *D*_xy_ among subsampled observations = 0.0111; Supplementary Fig. 2a, blue histogram) in comparison to *D*_xy_ between the outcrosser and the selfer when genomic windows were characterized by admixed HMM calls (average *D*_xy_ among subsampled observations = 0.0105; Figure 2A, red histogram). The distribution of the difference between *D*_xy_ calculated with admixed windows and *D*_xy_ calculated with control windows was greater than zero in the observed (empirical) data and centered on zero in the permuted data (Supplementary Fig. 2b). The observed distribution was significantly greater than the permuted data (p = 0.023; 97.7% of the difference in observed and permuted distributions was greater than zero; Supplementary Fig. 2c), suggesting that sequences called “admixed” by the HMM are truly the result of introgression from *xantiana*.

We found a similar pattern at site SM, although the effect of HMM call was marginally significant. *D*_xy_ between the selfer and outcrosser was greater for genomic windows characterized by conspecific HMM calls (average *D*_xy_ among subsampled observations = 0.0108; Supplementary Fig. 2b, blue histogram) than by those characterized by admixed HMM calls (average *D*_xy_ among subsampled observations = 0.0101; Supplementary Fig. 2a, red histogram). The distribution of the difference between *D*_xy_ calculated with admixed windows and *D*_xy_ calculated with control windows was greater than zero in the observed (empirical) data and centered on zero in the permuted data (Supplementary Fig. 2e), although there was marginally more overlap between distributions than for the S22 data (Supplementary Fig. 2b). Consequently, there was a marginally significant effect – sequences called “admixed” by the HMM had lower *D*_xy_ with the outcrosser than those called “unadmixed” (i.e., control in Supplementary Fig. 2d; p = 0.069, 93.1% of the difference in observed and permuted distributions was greater than zero; Supplementary Fig. 2f).

#### **Alternative methods to quantifying admixture proportions**

##### ***Methods***

In the main text, we quantify individual-level admixture proportions using a HMM, which infers introgression across the genome of an individual. To corroborate these results, we use two other methods of quantifying admixture proportions, both of which estimate admixture at the population level.

*D_xy_-based estimates of the admixture proportion*

The first method is a *D*_xy_-based calculation of admixture proportion. To simplify analysis, we consider the case of unidirectional introgression, which can be enforced by considering appropriate population comparisons (i.e. by making use of populations which could are unlikely to have previously experienced admixture). To do so we consider a population’s genomic variation as a combination of introgressed and non-introgressed ancestry in proportions α and 1 – α, respectively. As such, α of the genome is introgressed with “interspecific divergence” actually reflecting variation within the donor population ( π*, where the * denotes that this is an idealized donor population that has not experienced introgression), and 1 – α of the genome is non-introgressed with “interspecific divergence” in these regions reflecting divergence between species (hereafter *D*_xy_*, with the * noting we are considering divergence in the absence of introgression). Observed sequence divergence between any two populations, *D*_xy_, equals:

*D*_xy_ = α π* + (1 – α) *D*_xy_*	[Eq 1a]

We can rearrange this this equation to solve for α:

α = (*D*_xy_ * – *D*_xy_) / (*D*_xy_ * – π*)	[Eq 1b]

We have an estimate of *D*_xy_ from our sequence data, but do not know the values of the theoretical quantities *D*_xy_* (expected divergence between populations in the absence of introgression) and π* (expected intraspecific differentiation between the introgressed material and its source species). However, we can use our population sampling to find reasonable stand-ins (from allopatric *xantiana* (LCW and DLE) and *parviflora* (YC) populations near the region of sympatry for the values (see below).

We show in the main text that divergence between allopatric *xantiana* and *parviflora* is somewhat variable – suggesting a complex history of divergence, rather than a simple population split. We therefore provide two estimates of *D*_xy_*: (1) Median divergence between *parviflora* and the *xantiana* LCW population – the allopatric sample with the lowest sequence divergence from *parviflora*. In fact, *D*_xy_ between *xantiana* LCW and *parviflora* is less than that of some sympatric *xantiana* populations and *parviflora*, potentially suggesting gene flow into an ancestor of LCW, introgression from a further diverged *xantiana* population into some sympatric xantiana populations and/or some other complicated history of divergence. (2) Median divergence between *parviflora* and the *xantiana* DLE population – the allopatric population nearest the zone of sympatry, with *D*_xy_ to *parviflora* that exceeds that of all sympatric *xantiana* populations. We note that because *D*_xy_ between some allopatric *xantiana* populations (most notably THPO and CF) and *parviflora* exceeds *D*_xy_*, we estimate negative values of introgression in some cases. These negative values reflect the complex history of divergence in this group – consistent with either introgression from *parviflora* into an historical *xantiana* population which gave rise to some of the currently allopatric populations, or a complicated history of populations splitting (e.g. the selfer, *parviflora*, ‘budded’ from within the ancestral range of the outcrosser, *xantiana*, or a history of gene flow between some xantiana populations and some more distant and unsampled *Clarkia* species).

Our estimate also required a value of π*, intraspecific diversity within the donor. We simply assert π* to be 0.4% – roughly the level of divergence between the nearest allopatric *parviflora* (P_YC) and sympatric *parviflora* samples, although we provide a few examples of how changing the value impacts our estimated admixture proportion.

Finally, we note that because levels of interspecific divergence are relatively similar to levels of divergence between *xantiana* populations (*D*_xy_* ≈ π*), using this approach to estimate the extent of introgression of *xantiana* DNA into *parviflora* led to unstable and unreliable values – as the denominator was approximately zero – which we do not report here.

*f_d_ -based estimates of the admixture proportion*

We also provide *f_d_* -based estimates , of the fraction of the admixture proportion (Martin et al. 2015). Similar to ABBA-BABA analyses, this method uses three populations and an outgroup, (((P1, P2), P3,O). The f_d_ statistic is calculated by taking the ratio of the difference in sums of ABBA and BABA in the sample relative to that under a scenario of complete introgression. We used a combination of scripts from Martin’s Github tutorial “ABBA BABA statistics using genome wide SNP data (<https://github.com/simonhmartin/tutorials/blob/master/ABBA_BABA_whole_genome/README.md>) and from Martin et al. (2015) to calculate genome-wide *f_d_ .* We used the block jack-knifing script from Martin’s Github tutorial to calculate 95% confidence intervals on *f_d_* values.

To calculate admixture proportions into each of the contact zone populations, we used the same four-taxon contrasts that we used in ABBA-BABA analyses. To quantify admixture into *xantiana*, we used the (P1 = close allopatric xantiana [X_DLE], P2 = sympatric xantiana, P3 = close allopatric parviflora [P_FAY], P4 = outgroup [*Clarkia unguiculata*]) quartet structure, using a separate sympatric *xantiana* population as p2 in each analysis. To quantify admixture into *parviflora*, we used the (P1 = close allopatric parviflora [P_FAY], P2 = sympatric parviflora, P3 = close allopatric xantiana [X_DLE], P4 = outgroup [*Clarkia unguiculata*]) quartet structure, using a separate sympatric *xantiana* population as P2 in each analysis.

We also explore the effect of changing the non-focal taxa, in particular the P1 and P3 taxa for each contrast.

##### ***Results***

###### *Dxy-based calculations of admixture proportions*

We used patterns of divergence within and between populations and subspecies to estimate the extent of introgression from the selfer, *parviflora*, into the outcrosser, *xantiana*. We infer the highest level of *parviflora* ancestry in northern *xantiana* populations – roughly 30% in GC, roughly 13% in S22 and roughly 10% in SAW (Supplementary Fig. 3). Consistent with our D_xy_-based and ABBA-BABA tests for the presence of introgression, we find substantially less admixture in southern sympatric xantiana populations – approximately 3% in SM and S7, and negligible levels of introgression in GRW (Supplementary Fig. 3).

We also find negative admixture proportions in the geographically distant xantiana populations – CF, OC, and THPO (Supplementary Fig. 3). These negative estimates of the admixture proportion are consistent with other evidence of introgression into our allopatric populations and/or a speciation history more complex than expected from a simple population split.

While these D_xy_-based estimates of admixture proportions differ slightly based on whether LCWW or DLE is used as the reference *xantiana* population (linetypes in Supplementary Fig. 3), they can vary much more depending on the assumed divergence between introgressed *parviflora* ancestry and a reference *parviflora* source, π* (line colors, Supplementary Fig. 3). If we change our assumed value of π* from 0.4% to 0.7% we estimate an admixture proportion of nearly 50% in GC, while decreasing π* to 0.2% decreases the estimated admixture proportion to 25% in GC.

###### *f_d_-based calculations of admixture proportions*

The *f_d_* statistic estimated non-zero admixture proportions in sympatric *xantiana* populations but not in sympatric *parviflora* populations (Supplementary Fig. 4a-b and Supplementary Table 5).

In *xantiana* (Supplementary Fig. 4a)*,* the highest admixture proportion was estimated in the GC population (0.246, 95% CI: 0.228 - 0.63), followed by S22 (0.078, 95% CI: 0.067 - 0.089), SAW (0.050, 95% CI: 0.040 - 0.061), and SM (0.026, 95% CI: 0.015 - 0.036). The *f_d_* estimate was not statistically different from zero in the *xantiana* S7 (0.008, 95% CI: -0.001 - 0.016) and GRW (-0.001, 95% CI: -0.011 - 0.008) populations, consistent with low levels of introgression detected in these populations with the methods above.

Although all such estimates depend on the other populations used in the four-taxon comparison, our results (presented in Supplementary Fig. 4a,b with the ABBA-BABA comparison used in the main text) are quite consistent regardless of which allopatric *xantiana* population is used as the P1 taxon, with one small caveat (Supplementary Fig. 4c). When very distant *xantiana* populations are used as the outgroup (e.g., population OC, Supplementary Fig. 4c), we further increase our estimated admixture proportion in sympatric *xantiana* samples. This again could reflect either a budding origin of *parviflora*, gene flow from an unsampled *xantiana* population into the CF and OC populations and/or historical introgression of *parviflora* ancestry into ancestors of populations that are currently allopatric.

###### *Comparison of f_d_ and D_xy_-based estimates with HMM estimates of admixture proportion*

We compare the three methods for estimating admixture proportion in *xantiana* contact zones. Because the f_d_ and D_xy_-based estimates are underpowered to detect introgression into *parviflora*, we do not do this comparison for the *parviflora* contact zones. The three methods’ estimates of admixture proportion across sympatric *xantiana* populations are quite similar in rank order - i.e., the *xantiana* population at site GC always has the highest admixture proportion, followed by the *xantiana* population at site S22 (Extended Data Fig. 7).

### **Supplementary Figures**

#### **Supplementary Fig. 1:** A diagram of our approach to test the HMM-based inference of introgression of *xantiana* (outcrosser) ancestry into *parviflora* (selfer). See Supplementary text ‘Validation of introgression into the selfer’ for details.

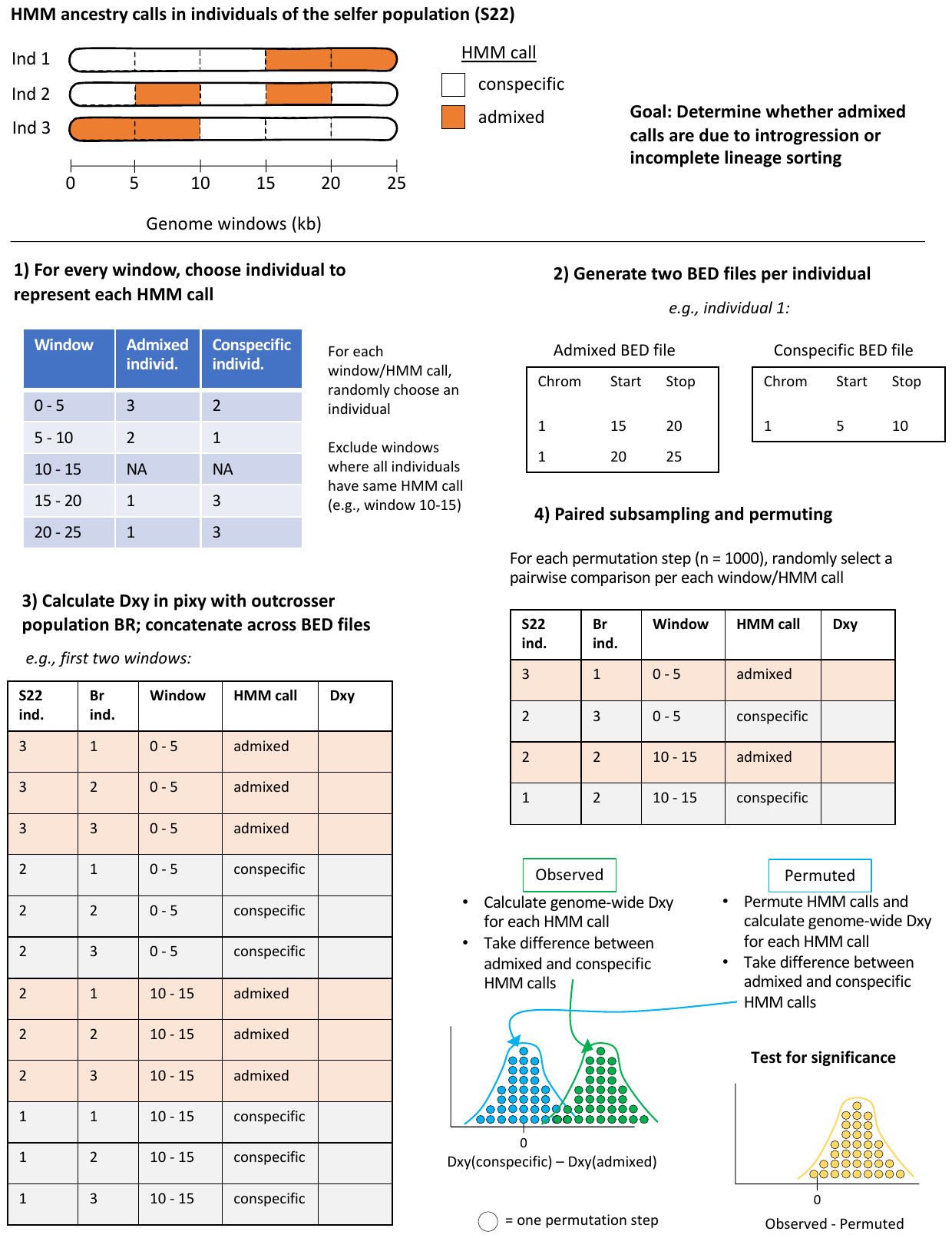

**Figure 2: Validation of HMM provides evidence for true introgression into the selfer, *parviflora*, at contact zones S22 and SM.** We find support for the HMM’s inference of introgression from the outcrosser, *xantiana*, into the selfer, *parviflora*, at both site S22 (A, B, C) and SM (D, E, F). (A,D) Estimates of *D*_xy_ between the selfer and outcrossing population BR when the genome is a mosaic of individuals with admixed HMM calls versus conspecific/unadmixed/control HMM calls. If introgression is the result of admixed HMM calls, we expect *D*_xy_ with the outcrosser to be lower for admixed calls than control calls. These are distributions, instead of point estimates, from the subsampling of data to remove the pseudoreplication introduced by calculating *D*_xy_ for a given genomic window/HMM call with 5 individuals from the outcrossing population BR. (B, E) Distributions of the observed and permuted difference between *D*_xy_ of admixed and control calls. (C, F) We test the null hypothesis that *D*_xy_ of the genomic regions with admixed HMM calls are less than that with unadmixed HMM calls by subtracting the permuted distribution from the observed distribution of this difference, and quantifying the percent of this derived distribution that is greater than zero.

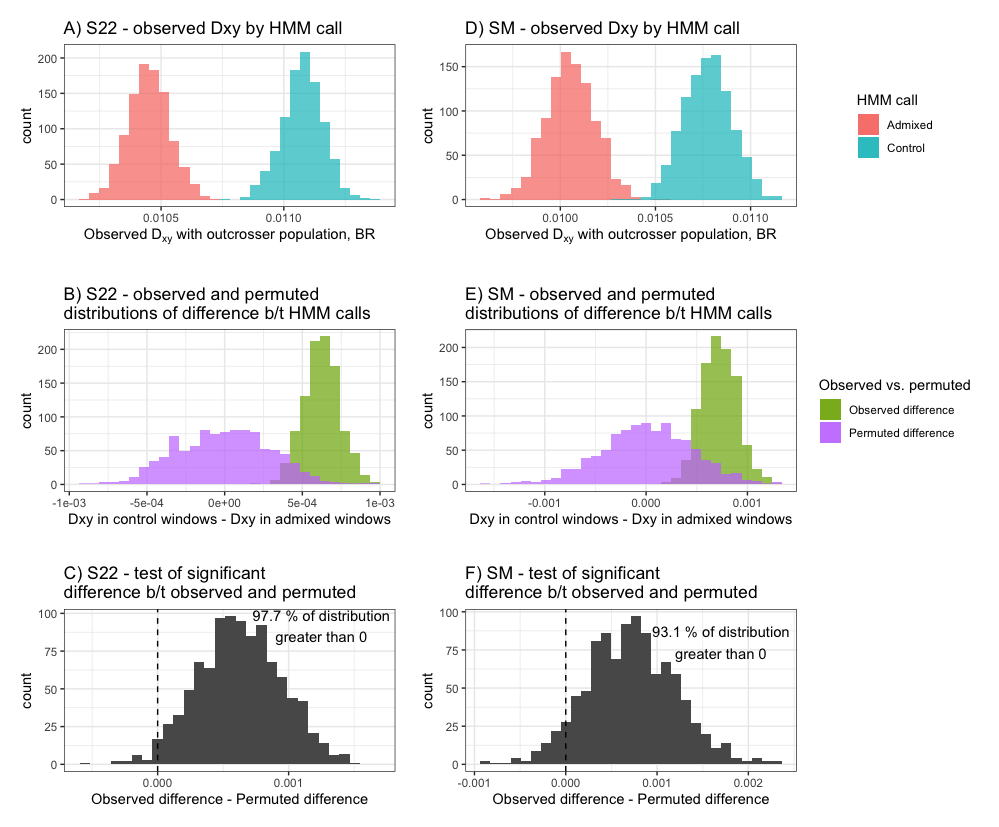

**Supplementary Fig. 3: D_xy_-based calculations of admixture proportion (α) infer non-zero admixture proportions in five sympatric populations (S7, SM, SAW, S22, GC).** X-axis is the population in which admixture is being inferred. We visualize the effects of two variables on admixture proportion: the reference *xantiana* population used to calculate the expected interspecific D_xy_ with *parviflora* (D_xy_* in Equation 1a), and the value used for the expected intraspecific diversity within the donor subspecies, *parviflora* (π*, Equation 1a).

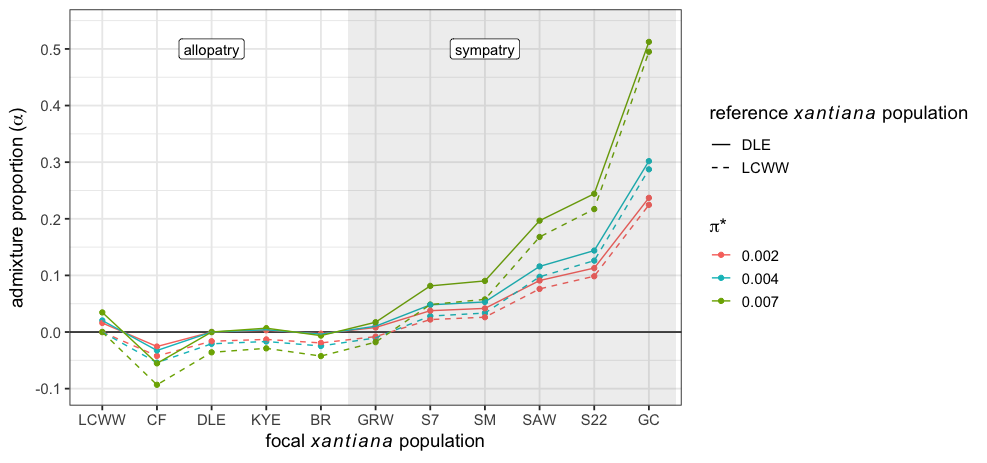

**Supplementary** **Fig. 4: *f_d_* estimates of admixture proportion.** (A) Estimates of admixture proportions in sympatric *xantiana* and (B) *parviflora* populations using the same four-taxon quartets as in the ABBA-BABA analyses in the main text. (C) Because this method only estimates non-zero admixture proportions in *xantiana*, we explore how the allopatric *xantiana* population (P1 in the four-taxon quartet) affects estimates of *f_d_*. Points and error bars represent the mean and 95% confidence intervals, respectively, of block jack-knifed estimates.

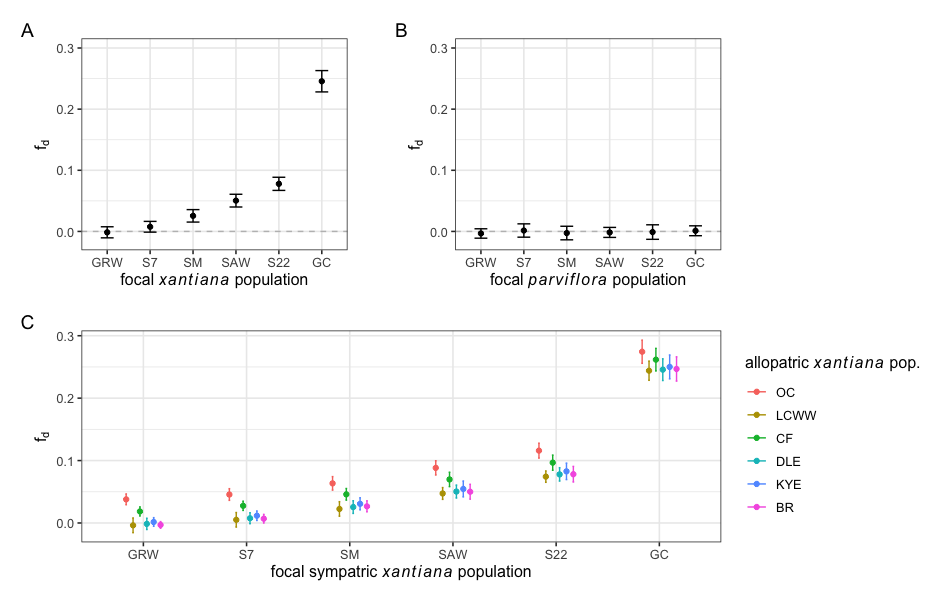

#### **Supplementary Tables**

**Supplementary Table 1: Population sampling of *Clarkia xantiana* subspecies and four outgroup species.** “Patry” refers to whether populations were allopatric or sympatry (i.e., at contact zones).

| Species | Patry | Population | Population Abbreviation | Individuals sampled | Latitude (WGS84) | Longitude (WGS84) | Elevation (m) |
| --- | --- | --- | --- | --- | --- | --- | --- |
| *Clarkia xantiana*  *ssp. parviflora*   (selfer) | allopatry | Valyermo | VALY | 2 | 34.438339 | -117.841573 | 1173 |
|  |  | Short Canyon | SRTCAN | 2 | 35.7136075 | -117.9175 | 1149 |
|  |  | Long Valley | LV | 5 | 35.8166587 | -118.08334 | 1685 |
|  |  | 20p | 20P | 6 | 35.6997804 | -118.06284 | 1413 |
|  |  | 100p | 100P | 6 | 35.7329102 | -118.08345 | 1246 |
|  |  | Piute Mountain | 48P | 6 | 35.4679364 | -118.22417 | 1400 |
|  |  | Kelso Valley | KV | 5 | 35.6158761 | -118.25255 | 913 |
|  |  | White Blanket | WB | 6 | 35.6999916 | -118.21667 | 887 |
|  |  | Fay Ranch | FAY | 6 | 35.7315606 | -118.31062 | 1227 |
|  |  | Yankee Canyon | YC | 6 | 35.6499952 | -118.43334 | 923 |
|  | sympatry | Green Rock W | GRW | 8 | 35.5973049 | -118.50963 | 739 |
|  |  | Site 7 | S7 | 8 | 35.5943338 | -118.50715 | 807 |
|  |  | Squirrel Mtn | SM | 9 | 35.6020115 | -118.41679 | 1218 |
|  |  | Sawmill Rd E | SAW | 9 | 35.6795127 | -118.47434 | 913 |
|  |  | Golf Course | GC | 8 | 35.7266285 | -118.43094 | 803 |
|  |  | Site 22 | S22 | 9 | 35.8377624 | -118.44963 | 929 |
| *Clarkia xantiana*  *ssp. xantiana*  (outcrosser) | allopatry | Three Points | THPO | 5 | 34.7377213 | -118.59963 | 1084 |
|  |  | Oiler Canyon | OC | 5 | 35.339932 | -118.58165 | 845 |
|  |  | Lucas Creek W | LCW | 6 | 35.4809755 | -118.71036 | 517 |
|  |  | Cow Flat | CF | 4 | 35.5296301 | -118.65358 | 661 |
|  |  | Delonegha E | DLE | 4 | 35.5464294 | -118.61676 | 670 |
|  |  | Keyesville E | KYE | 5 | 35.622377 | -118.51466 | 980 |
|  |  | Borel Road | BR | 6 | 35.5883736 | -118.51728 | 804 |
|  | sympatry | Green Rock W | GRW | 8 | 35.5973049 | -118.50963 | 739 |
|  |  | Site 7 | S7 | 8 | 35.5943338 | -118.50715 | 807 |
|  |  | Squirrel Mtn | SM | 8 | 35.6020115 | -118.41679 | 1218 |
|  |  | Sawmill Rd E | SAW | 9 | 35.6795127 | -118.47434 | 913 |
|  |  | Golf Course | GC | 9 | 35.7266285 | -118.43094 | 803 |
|  |  | Site 22 | S22 | 9 | 35.8377624 | -118.44963 | 929 |
| *Clarkia unguiculata* |  | Democrat | DEM | 3 | 35.5265165 | -118.62799 | 828 |
| *Clarkia cylindrica* |  | Democrat | DEM | 2 | 35.5265165 | -118.62799 | 828 |
| *Clarkia speciosa* |  | Black Gulch | BG | 3 | 35.5911972 | -118.52785 | 727 |
| *Clarkia exilis* |  | Old Kern Canyon Rd | OKC | 3 | 35.5636912 | -118.56714 | 815 |

**Supplementary Table 2: ABBA-BABA detects introgression into the outcrosser but not the selfer.** We tested for introgression into each taxon at each of the six contact zone sites. For tests of introgression into *xantiana*, the outcrosser, we set taxon P2 as the focal contact zone site. The taxa for P1 (allopatric outcrosser), P3 (allopatric selfer) and P4 (outgroup) remained constant. Taxa are indicated by “Subspecies (X or P)_SITE”. We present the number of ABBA (introgression from P3 to P2), BABA (introgression from P3 to P1) and BBAA (species tree) trees for each test; we predicted an excess of ABBA trees and a positive D statistic if introgression is occurring in contact zones. The 95% confidence intervals are based on 1000 bootstrap replicates. Significant D statistics are bolded.

|  | **Taxa used in each test** | | | | **Number of sites** | | | | **D-statistic** | | | |  |
| --- | --- | --- | --- | --- | --- | --- | --- | --- | --- | --- | --- | --- | --- |
|  | **p1** | **p2** | **p3** | **p4** | **Total** | **ABBA** | **BABA** | **BBAA** | **D** | **std error** | **lower 95% CI** | **higher 95% CI** | **95% CI overlaps zero** |
| Tests for introgression into focal  **xantiana (outcrosser)**  sympatric population (p2) | X_DLE | X_S22 | P_FAY | UNG | 6307.06 | 2246.45 | 1675.80 | 2384.81 | **0.145** | 0.007 | 0.132 | 0.158 | No |
|  |  | X_GC |  |  | 6537.64 | 3043.51 | 1455.40 | 2038.74 | **0.353** | 0.006 | 0.342 | 0.365 | No |
|  |  | X_SAW |  |  | 6301.78 | 2024.97 | 1666.71 | 2610.10 | **0.097** | 0.007 | 0.084 | 0.111 | No |
|  |  | X_SM |  |  | 6464.64 | 1927.91 | 1733.62 | 2803.11 | **0.053** | 0.007 | 0.038 | 0.066 | No |
|  |  | X_S7 |  |  | 6450.27 | 1814.38 | 1751.32 | 2884.57 | **0.018** | 0.008 | 0.003 | 0.034 | No |
|  |  | X_GRW |  |  | 6482.88 | 1770.97 | 1784.24 | 2927.67 | -0.004 | 0.007 | -0.018 | 0.011 | Yes |
| Tests for introgression into focal  **parviflora (selfer)**  sympatric population (p2) | P_FAY | P_S22 | X_DLE | UNG | 6751.69 | 877.00 | 874.90 | 4999.79 | 0.001 | 0.012 | -0.023 | 0.026 | Yes |
|  |  | P_GC |  |  | 6936.31 | 648.68 | 638.80 | 5648.84 | 0.008 | 0.015 | -0.021 | 0.037 | Yes |
|  |  | P_SAW |  |  | 6855.15 | 704.15 | 701.12 | 5449.88 | 0.002 | 0.013 | -0.023 | 0.026 | Yes |
|  |  | P_SM |  |  | 6811.66 | 800.08 | 814.19 | 5197.39 | -0.009 | 0.013 | -0.034 | 0.016 | Yes |
|  |  | P_S7 |  |  | 6872.39 | 681.48 | 681.05 | 5509.86 | 0.000 | 0.015 | -0.031 | 0.031 | Yes |
|  |  | P_GRW |  |  | 6966.36 | 591.70 | 605.67 | 5768.99 | -0.012 | 0.016 | -0.042 | 0.023 | Yes |

**Supplementary Table 3: The magnitude of introgression varies between subspecies and across contact zone sites.** Average individual admixture proportions (2*standard error) inferred from the HMM.

|  | ***S22*** | ***GC*** | ***SAW*** | ***SM*** | ***S7*** | ***GRW*** |
| --- | --- | --- | --- | --- | --- | --- |
| ***xantiana* (outcrosser)** | 0.255 (0.018) | 0.348 (0.162) | 0.152 (0.008) | 0.065 (0.013) | 0.014 (0.004) | 0.013 (0.001) |
| ***parviflora* (selfer)** | 0.069 (0.019) | 0.012 (0.002) | 0.018 (0.003) | 0.038 (0.011) | 0.012 (0.001) | 0.009 (0.001) |

**Supplementary Table 4: Peak flowering time overlap between the outcrosser (*xantiana,* X) and the selfer (*parviflora,* P) at seven contact zones across two years, summarized from data in Eckart and Geber (1999).** Columns “X” and “P” give Julian dates (preceded by last two digits of year) for peak flowering time. “X-P” columns give the difference, in days, between peak flowering of *xantiana* and *parviflora*. The last column shows how much the difference between subspecies varies between the years. The year 1998 was an El Niño (dried) year and 1999 was a La Niña (wetter) year.

|  | **1998** | | | **1999** | | | **Difference between years in overlap** |
| --- | --- | --- | --- | --- | --- | --- | --- |
|  | **X** | **P** | **X - P** | **X** | **P** | **X - P** |  |
| ***Site 22*** | 98164 | 98137 | 27 | 99148 | 99138 | 10 | 17 |
| ***Site 5 (Chico Flat)*** | 98154 | 98135 | 19 | 99146 | 99132 | 14 | 5 |
| ***Site 77/78 (Sawmill)*** | 98165 | 98138 | 27 | 99152 | 99131 | 21 | 6 |
| ***Site 74 (Bodfish Cyn)*** | 98169 | 98142 | 27 | 99151 | 99137 | 14 | 13 |
| ***Site 70 (Erskine Crk)*** | 98161 | 98135 | 26 | 99152 | 99128 | 24 | 2 |
| ***Site 7*** | 98165 | 98150 | 15 | 99150 | 99139 | 11 | 4 |
| ***Site 9*** | 98156 | 98142 | 14 | 99148 | 99137 | 11 | 3 |
| Mean |  |  | 20.5 |  |  | 15 |  |

Eckhart, VM and MA Geber. "Character variation and geographic distribution of Clarkia xantiana A. Gray (Onagraceae): flowers and phenology distinguish two subspecies." *Madroño* (1999): 117-125.^17^

**Supplementary Table 5: Quantification of admixture proportion within each taxon at each contact zone (P2) with the *f_d_* statistic.** Block jackknifing was used to calculate a distribution of *f*_d_ values (admixture proportion), from which we calculated the mean and the 95% confidence interval. *f_d_* values that are significantly greater than zero are bolded.

|  | **P1** | **P2** | **P3** | **P4** | **mean *f_d_*** | **95% Confidence interval** | |
| --- | --- | --- | --- | --- | --- | --- | --- |
|  |  |  |  |  |  | **lower** | **upper** |
| introgression into *xantiana* (outcrosser) | X_DLE | X_S22 | P_FAY | UNG | **0.0778** | 0.0671 | 0.0886 |
|  |  | X_GC |  |  | **0.2456** | 0.2282 | 0.2631 |
|  |  | X_SAW |  |  | **0.0504** | 0.0399 | 0.0608 |
|  |  | X_SM |  |  | **0.0255** | 0.0153 | 0.0357 |
|  |  | X_S7 |  |  | 0.0076 | -0.0012 | 0.0164 |
|  |  | X_GRW |  |  | -0.0014 | -0.0105 | 0.0076 |
| introgression into *parviflora* (selfer) | P_FAY | P_S22 | X_DLE | UNG | -0.0010 | -0.0129 | 0.0108 |
|  |  | P_GC |  |  | 0.0011 | -0.0070 | 0.0092 |
|  |  | P_SAW |  |  | -0.0016 | -0.0098 | 0.0066 |
|  |  | P_SM |  |  | -0.0026 | -0.0137 | 0.0085 |
|  |  | P_S7 |  |  | 0.0015 | -0.0094 | 0.0124 |
|  |  | P_GRW |  |  | -0.0034 | -0.0111 | 0.0043 |
