## Extended Data for "Mating system and environment predict the direction and extent of introgression between incipient *Clarkia* species"

**Extended Data Fig. 1 Genetic diversity metrics within subspecies is consistent with mating system shifts.**

The selfer, *parviflora*, has A) lower π within populations, B) lower *D*_xy_ (absolute divergence) between pairs of population, and C) higher *F*_ST_ (relative divergence) between pairs of populations than the outcrosser, *xantiana*. All diversity metrics were calculated at 4-fold degenerate coding region sites.

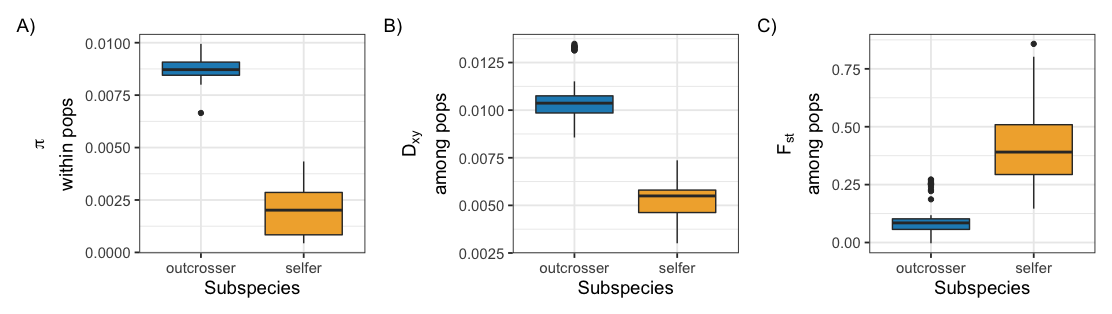

**Extended Data Fig. 2 Genetic structure of subspecies largely reflects geography.**

A) Maps of sampling locations. Left panel shows all populations, including geographic outliers THPO in *xantiana* and VALY in *parviflora*. Right panel has populations within subspecies shaded by geography, from light hues in allopatry to dark hues in sympatry. Shading matches that in the Genetic PCAs (B-F). PCAs were calculated with 4-fold degenerate SNPs from coding sequences. B-D) *Xantiana* (outcrosser) genetic PCAs. Because the farthest allopatric populations THPO and OC each dominate PC space, we show PCAs calculated with THPO, and without THPO and OC. E-F) *Parviflora* (selfer) genetic PCAs. PCA was calculated with and without the geographic outlier VALY.

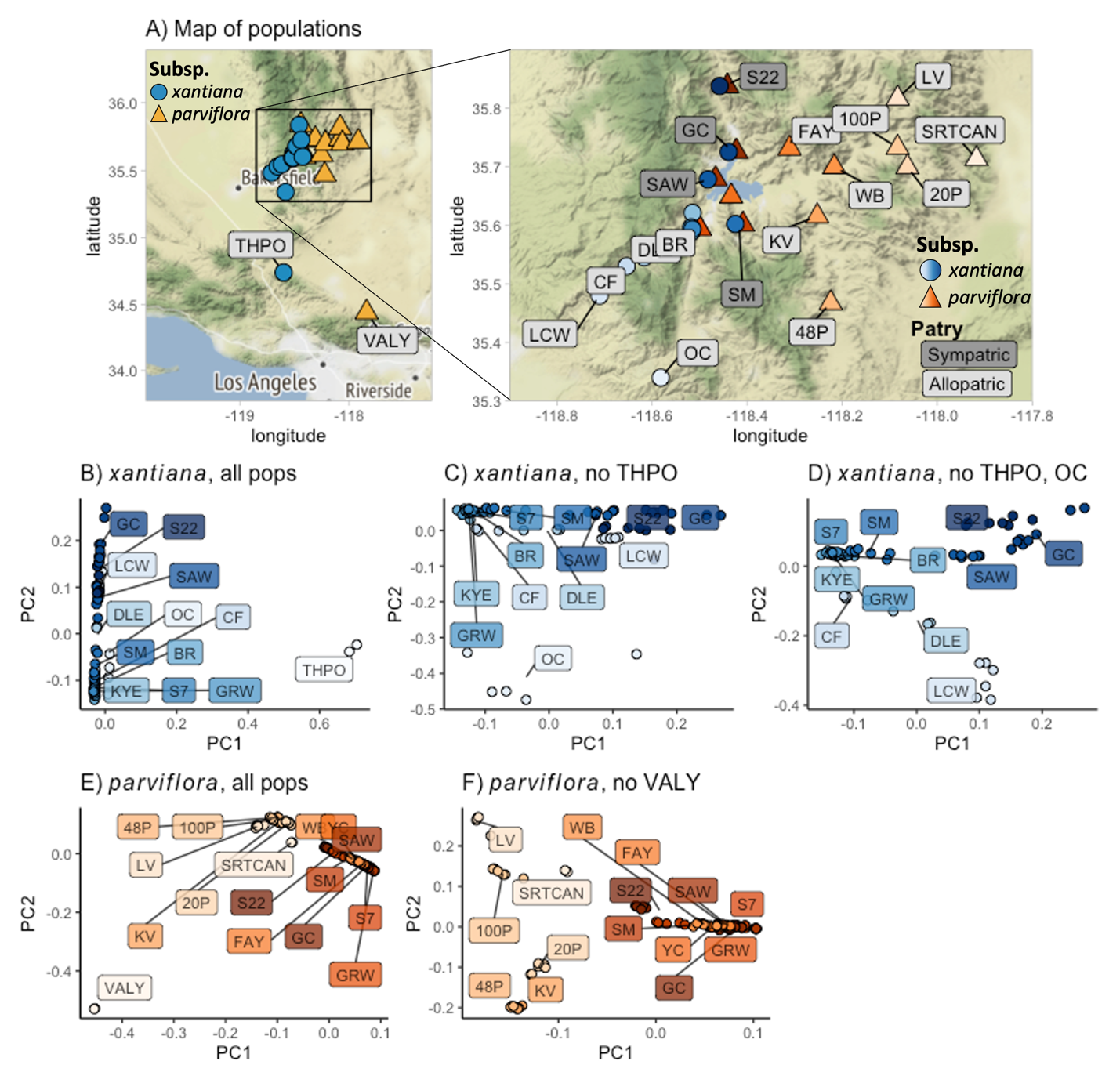

**Extended Data Fig. 3 ABBA-BABA tests of independent introgression across the contact zone.**

We used pairwise ABBA-BABA tests to determine whether signals of introgression across the contact zones are the result of ancestral or independent, local introgression. Panels A-C cartoon the four-taxa tree topologies that can arise in our ABBA-BABA tests. For a given pairwise combination, taxa can group by subspecies (A, species tree), or by two topologies discordant with the species tree: by site (B, indicates local introgression) or by neither subspecies nor site (C, indicates incomplete lineage sorting (ILS) of ancestral variation and/or introgression that occurred before subspecies split into the two sites). A significant excess of ABBA vs BABA trees is evidence for local introgression in one or both sites. (D) Heatmap of ABBA-BABA results between pairwise site combinations. Colors reflect D-statistics and asterisks show significant tests.

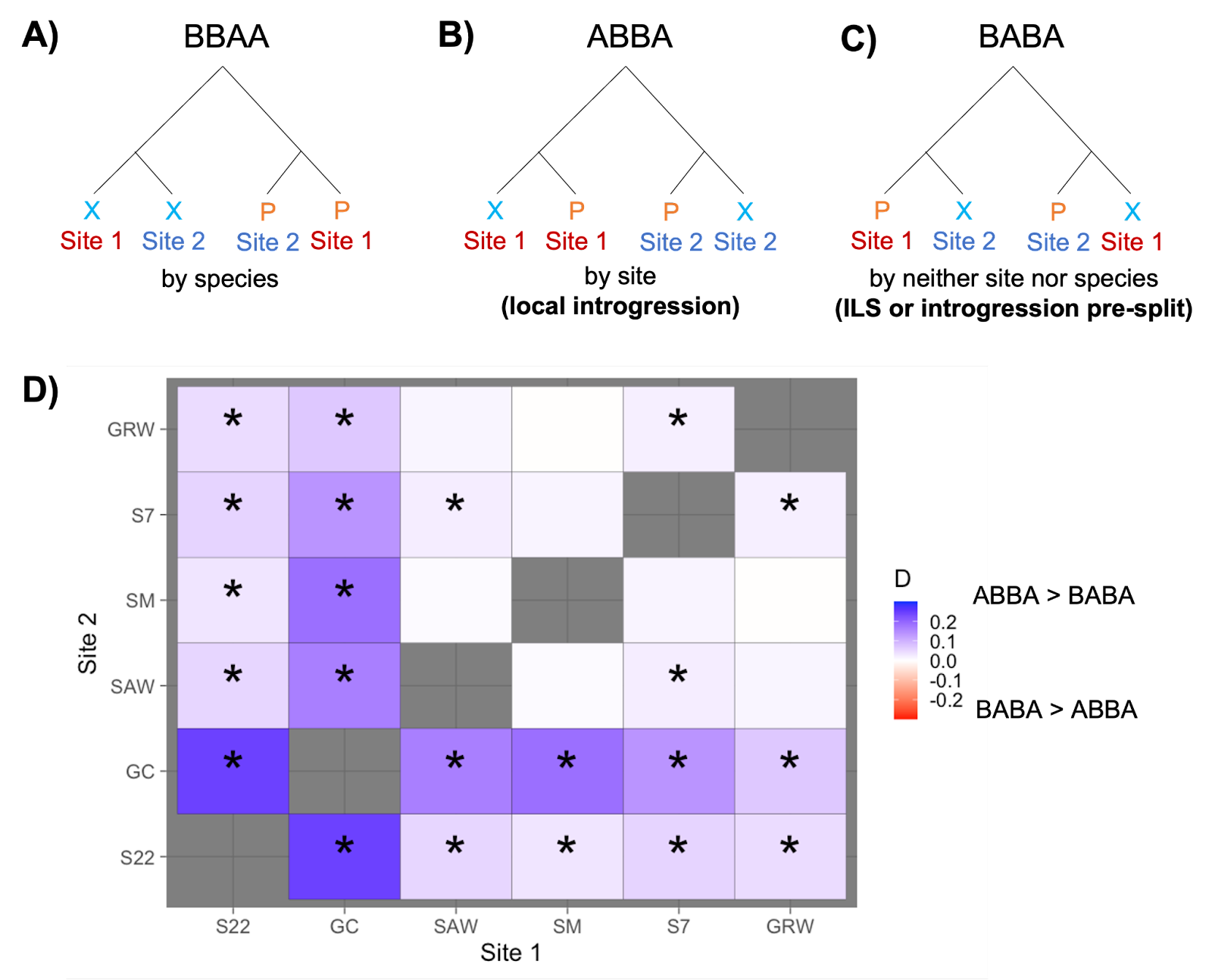

**Extended Data Fig. 4 Full chloroplast phylogeny of *xantiana* (blue), *parviflora* (orange), and outgroup species (black).** For *xantiana* and *parviflora* samples, labels are formatted as “Subspecies_population_individual”. Bootstrap labels for branches supporting outgroup species relationships, monophyly of *C. xantiana* as a whole, and monophyly of each subspecies are shown. The branch supporting monophyletic *xantiana* has a bootstrap value of 95%, whereas monophyly of the *parviflora* (plus some *xantiana* individuals) is more poorly supported (51% bootstrap).

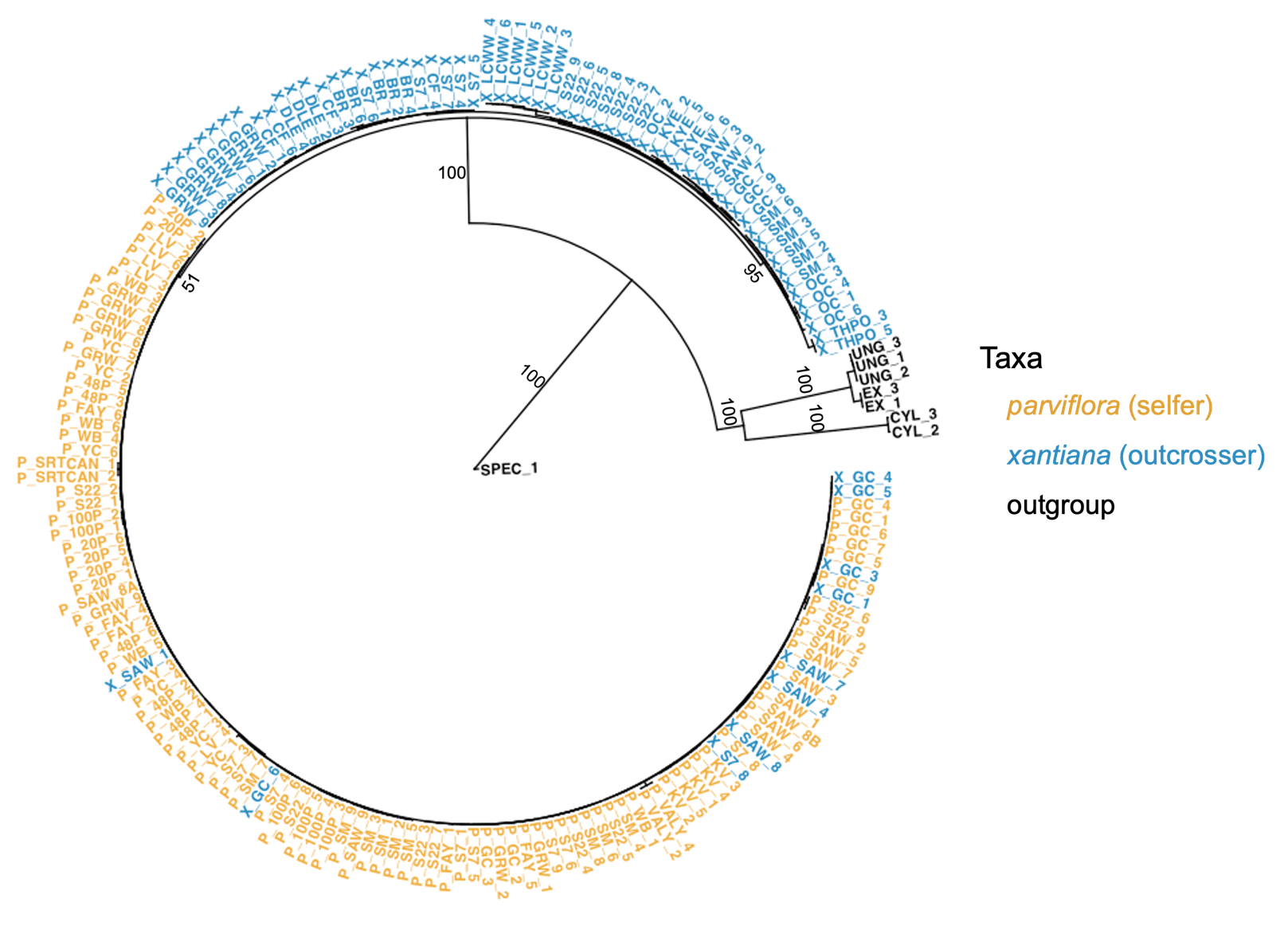

**Extended Data Fig. 5 Cladogram of the *parviflora* clade, highlighting the 10 *xantiana* individuals with captured chloroplasts.** For *xantiana* and *parviflora* samples, labels are formatted as “Subspecies_population_individual”. Bootstrap values greater than 75% are depicted.

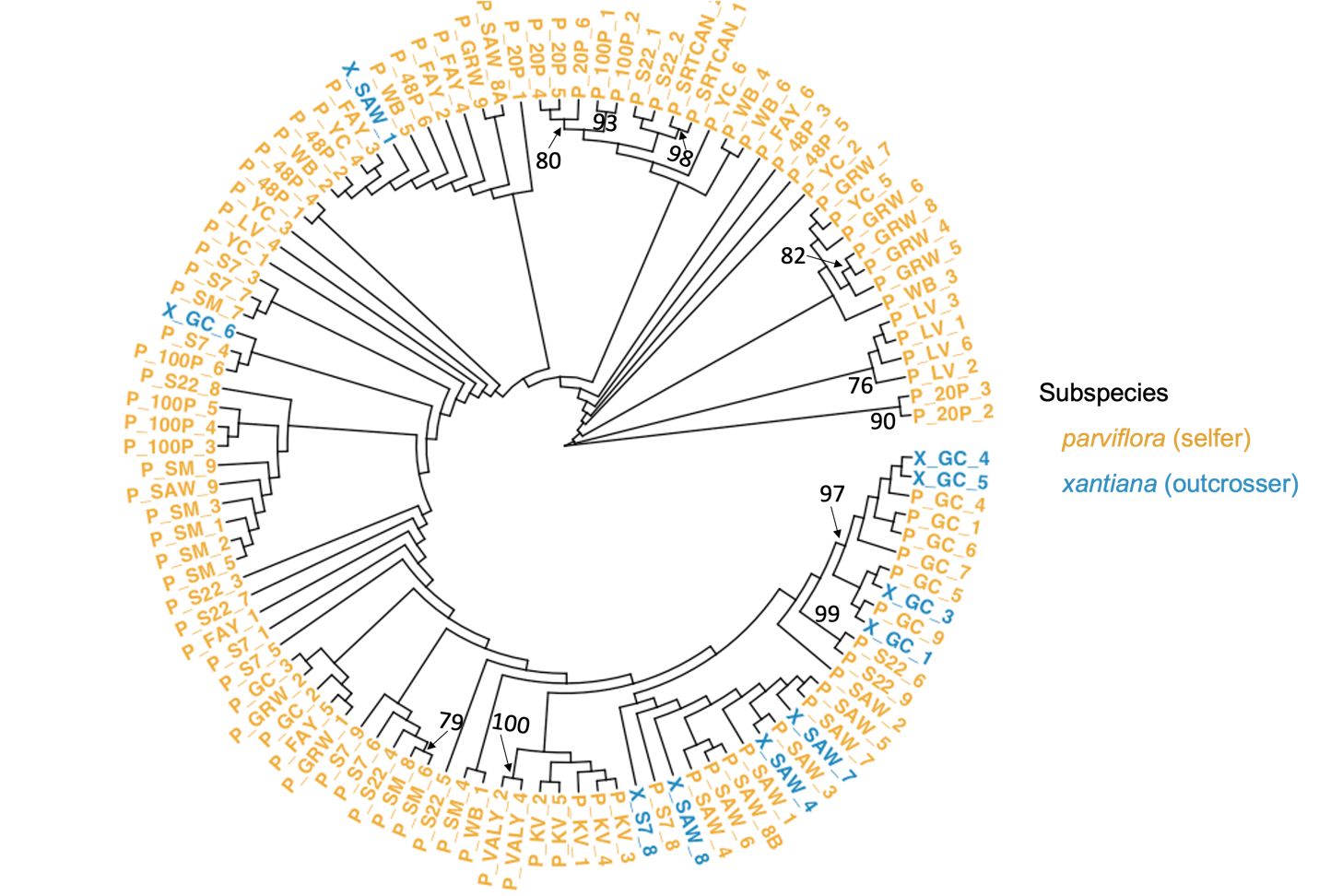

**Extended Data Fig. S6 Patterns of diversity within (pi) and between (D_xy_) sympatric *parviflora* populations provide evidence for introgression into *parviflora* at contact sites SM and S22.** In conjunction with HMM results, which find the highest admixture proportions at S22 and SM, patterns of diversity and divergence in these populations are consistent with introgression. We expect *parviflora* populations that experience introgression to be more divergent from other *parviflora* populations, and also to have higher within population diversity given that the admixed ancestry comes from a taxon with high diversity.

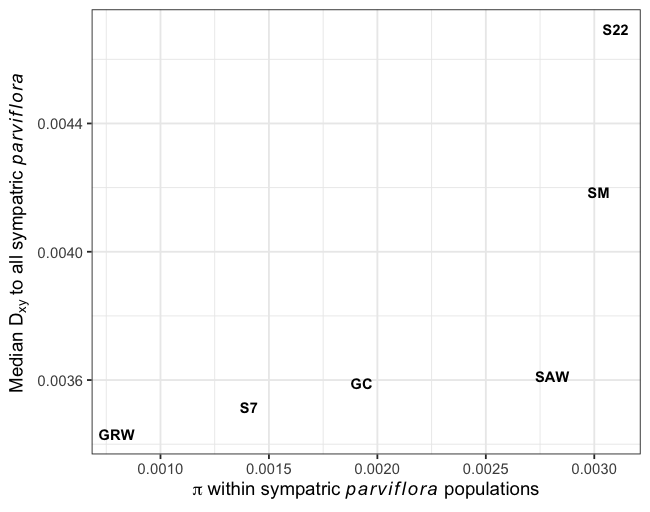

**Extended Data Fig. 7 Admixture proportions in *xantiana* estimated from different methods are highly correlated**. Each point represents the admixture proportion in the outcrosser, *xantiana*, for each of the six contact zones. We present the *D*_xy_ metric calculated with expected intraspecific diversity within *parviflora* (π*) as 0.004, the *f_d_* statistic calculated with population OC as the allopatric *xantiana* population (P1), and the HMM estimates calculated with either all sites or sites with high confidence ancestry calls. See Supplementary Text ‘Alternative methods to quantifying admixture proportions’ for details on analyses.

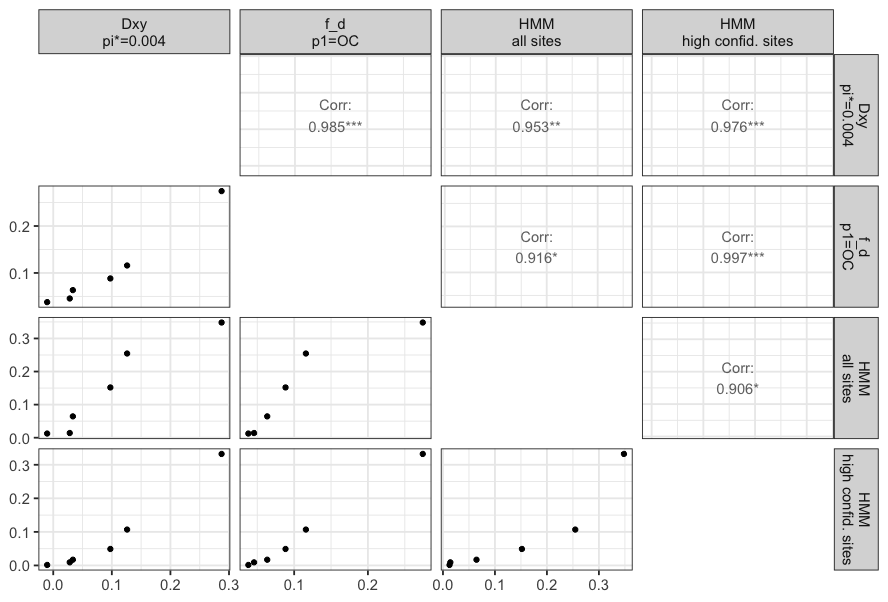
